# Stromal remodeling regulates dendritic cell abundance and activity in the tumor microenvironment

**DOI:** 10.1101/2021.11.10.467836

**Authors:** Athanasios Papadas, Gauri Deb, Adam Officer, Alexander Cicala, Chelsea Hope, Philip Emmerich, Joshua Wiesner, Adam Pagenkopf, Garrett Arauz, Varun Bansal, Kristina A. Matkowskyj, Dustin Deming, Katerina Politi, Scott I. Abrams, Olivier Harismendy, Fotis Asimakopoulos

## Abstract

Stimulatory dendritic cells (SDC), enriched within Batf3-DC (cDC1), engage in productive interactions with CD8+ effectors along tumor-stroma boundaries. The paradoxical accumulation of “poised” cross-presenting Batf3-DC within stromal sheets, distal to tumoral nests, is unlikely to simply reflect passive exclusion away from immunosuppressive tumor cores. Drawing parallels with embryonic morphogenesis, we hypothesized that invasive margin stromal remodeling may generate developmentally conserved cell-fate cues that regulate Batf3-DC behavior. We find that CD8+ T-cells massively infiltrate tumor matrices undergoing proteoglycan versican (VCAN) proteolysis, an essential organ-sculpting modification in development and adult tissue-plane forging. VCAN proteolysis releases a bioactive fragment (matrikine), versikine, that is necessary and sufficient for Batf3-DC accumulation. Versikine does not influence tumor-seeding pre-DC differentiation; rather, it orchestrates a distinctive activation program conferring exquisite sensitivity to DNA-sensing, coupled with survival support from atypical innate lymphoid cells. Thus, homeostatic signals from stroma invasion regulate SDC survival and activity to promote T-cell inflammation.

**HIGHLIGHTS:**

1. Tumor stroma remodeling generates cross-presenting DC survival and activation cues.
2. Stromal-activated Batf3-DC are hypersensitive to dsDNA-sensing.
3. Stromal signals promote atypical innate lymphoid cells (GM-CSF^hi^/ IFN*γ*^lo^).
4. T-cell repriming by stroma-licensed Batf3-DC may overcome exclusion at tumor margins.

## INTRODUCTION

Tumor antigen cross-presentation and CD8+ T cell effector priming by immunogenic Batf3- lineage DC (also known as type 1 conventional DC, cDC1) is integral to spontaneous and therapeutic anti-tumor immunity (Gajewski, 2015; Hildner et al., 2008). Innate sensing of tumors at the “elimination” phase of immunoediting depends on Batf3-DC (Binnewies et al., 2018; Broz et al., 2014; Salmon et al., 2016). In addition to effectively cross-priming CD8+ T cells recognizing tumor antigens in the lymph node, and re-priming CD8+ cytotoxic T lymphocytes (CTL) in the tumor bed, Batf3-DC regulate effector cell influx into the tumor microenvironment (TME) (Spranger et al., 2017). From a translational perspective, Batf3-DC are crucial for immunotherapy efficacy, including responses to vaccination strategies, immune checkpoint inhibitors (Oba et al., 2020; Salmon et al., 2016; Sanchez-Paulete et al., 2016) and engineered immune effector cells (e.g. CAR-T cells)(Kuhn et al., 2020).

Several key studies have shown that stimulatory Batf3-DC are excluded from interdigitating tumor nestlets and locate in areas of peritumoral stromal matrix (Bell et al., 1999; Broz et al., 2014; Hubert et al., 2020; Lavin et al., 2017; Mattiuz et al., 2021). However, the mechanisms that retain Batf3-DC at the peritumoral border remain poorly understood. To understand this paradoxical localization, we drew parallels with embryonic development where provisional matrix remodeling and plane-forging processes generate powerful cell fate-cues that regulate local cell behavior (Nandadasa et al., 2014). We hypothesized that homeostatic signals arising from the dynamic interfacing of the expanding tumor margin and the “defending” host stroma may regulate stimulatory Batf3-DC survival and activation *in situ*.

The tumor matrisome includes collagens, glycoproteins, and proteoglycans. Tumor matrix proteoglycans in particular have been implicated in nearly every hallmark of cancer. Arguably the most “versatile” member of the group (as its name suggests), versican (VCAN), possesses proven crucial roles in tumor growth, survival, invasion, metastasis and immune regulation, summarized in comprehensive review articles (Islam and Watanabe, 2020; Nandadasa et al., 2014; Papadas and Asimakopoulos, 2020; Wight et al., 2020). VCAN is essential for life and *Vcan*-null mice die in utero by embryonic day 10.5 due to defects along the anterior-posterior cardiac axis (Mjaatvedt et al., 1998). VCAN proteolysis by stromal fibroblast-derived ADAMTS (a disintegrin and metalloproteinase with thrombospondin motifs) proteases at the Glu^441^-Ala^442^ bond (V1 isoform enumeration) is essential for morphogenesis in the embryo, acting in part, through the specific activities of the released bioactive N-terminal fragment, versikine (McCulloch et al., 2009; Nandadasa et al., 2014; Timms and Maurice, 2020). Disruption of the Glu^441^-Ala^442^ proteolytic site that generates versikine leads to soft tissue syndactyly and other developmental abnormalities (Islam et al., 2020; Nandadasa et al., 2021). Aberrant VCAN proteolysis has also been associated with non-neoplastic structural tissue-plane pathologic changes in the adult (Fava et al., 2018).

Stromal VCAN proteolysis at the Glu^441^-Ala^442^ bond (V1 isoform enumeration) correlates with CD8+ T cell infiltration in both solid and hematopoietic tumors (Emmerich et al., 2020; Hope et al., 2017; Hope et al., 2014). Recombinant versikine protein elicited IRF8-dependent transcripts in myeloid cells *in vitro* in our hands (Hope et al., 2016), later confirmed by others (Han et al., 2020), and promoted generation of Batf3-DC from Flt3L-mobilized bone marrow (BM) *in vitro* (Hope et al., 2017). These activities of the proteolytic fragment versikine appear at odds with the immunosuppressive actions of parental non-proteolyzed VCAN (Tang et al., 2015). However, whether there is a linear connection between stromal matrix remodeling and adaptive anti-tumor immunity *in vivo* remains unknown. Here, we provide compelling evidence that connects tumor architecture dynamics with stimulatory DC abundance and function.

## RESULTS

### Essential roles of stromal remodeling signals in tumor Batf3-DC maintenance

In human epithelial cancers, peritumoral stromal sheets robustly accumulate matrix proteoglycans, including VCAN. VCAN sources in the tumor microenvironment include the stromal cells, immune infiltrating cells and in some cases, such as lung cancer, the tumor cells themselves (Papadas and Asimakopoulos, 2020). VCAN proteolytic processing however, is located primarily in the stroma due to the local activity of stromal fibroblast-derived ADAMTS versicanases (Hope et al., 2014). Using an immunohistochemistry (IHC)-validated antibody against DPEAAE, a neoepitope generated through VCAN proteolysis at Glu^441^-Ala^442^ (V1 isoform)(Fig. 1A), we observed DPEAAE signal in approximately 83% cases in a stromal distribution (Fig. 1B and Supp. Table S1). To determine the location of Batf3-DC relative to sites of stromal VCAN proteolysis, we performed IHC with an antibody detecting the Batf3-DC marker, XCR1 (Fig. 1B, inset). XCR1+ cells were localized in stromal sheets almost exclusively.

**Fig. 1.**
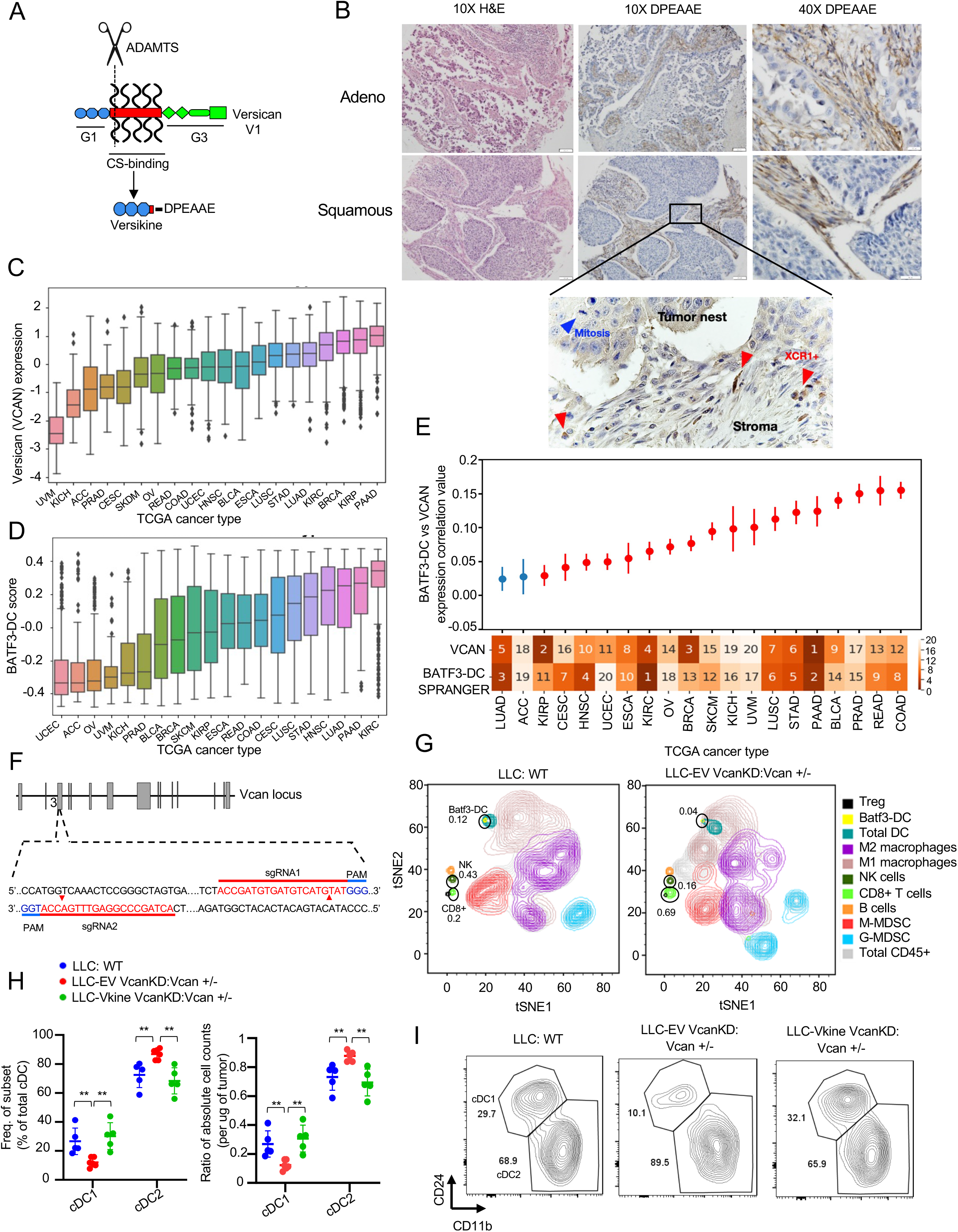
Essential roles of stromal remodeling signals in tumor Batf3-DC maintenance. A: Schematic showing versican (VCAN) functional domains and site-specific proteolysis site generating versikine (ADAMTS proteolytic cleavage= scissors). Versikine is generated through stromal fibroblast-derived ADAMTS-mediated proteolysis of the Glu^441^-Ala^442^ bond (V1 isoform enumeration), exposing a neoepitope (DPEAAE) that can be detected through a specific antibody. B: Stromal distribution of anti-DPEAAE immunohistochemical staining in human lung cancers. DPEAAE constitutes the C-terminus of versikine (chromogen: DAB, counterstain: hematoxylin). 10X objective: scalebar 50 μm, 40X objective: scalebar 20μm. Inset, XCR1+ IHC (chromogen: DAB, counterstain: hematoxylin). C: Distribution of VCAN expression across TCGA carcinomas, ordered on the horizontal axis by median VCAN expression. TCGA tumor code abbreviations are provided in gdc.cancer.gov. D: Distribution of measured Batf3-DC score (see Methods) across TCGA carcinomas, ordered on the horizontal axis by median measured Batf3-DC score. E: Levels of correlation between Batf3-DC score and VCAN expression across TCGA carcinomas. Ranked median of VCAN expression and measured Batf3-DC score shown across the X axis (1 highest, 20 lowest). Significant (q < 0.1) correlations after multiple hypothesis correction are colored in red. Error bars represent the standard error of the correlation coefficient measured using Python statsmodels. F: Generation of *Vcan*+/- mice through CRISPR-Cas9-based targeting of *Vcan* exon 3, encoding part of the G1 domain. Diagram showing sgRNA guide sequences, PAM motif and target sites. G: Mass cytometry demonstrating immune contexture in WT (LLC implanted into WT recipients- left) and VCAN-depleted tumor microenvironment (LLC^VcanKD^ tumor cells implanted in *Vcan*+/- recipients, right). H: Quantification of frequency (left) and absolute count ratios (cDC1/cDC1+cDC2 and cDC2/cDC1+cDC2) in WT, Vcan-depleted (Vcan+/-: LLC^VcanKD^-EV) and versikine-rescue (Vcan+/-: LLC^VcanKD^-Vkine) tumors. Data are represented as mean±SD. n=5 for each group. *p<0.05; **p<0.01;***p<0.001. I: Representative flow cytometry plots showing cDC1 and cDC2 frequency in WT, Vcan-depleted (Vcan+/-: LLC^VcanKD^-EV) and versikine-rescue (Vcan+/-: LLC^VcanKD^-Vkine) tumors (for gating strategy see Supp. Fig. 1E).

Matrikines (such as versikine) have been defined as “peptides liberated by partial proteolysis of extracellular matrix macromolecules which are able to regulate cell activities *not triggered by their full-size parent macromolecules”* (Gaggar and Weathington, 2016). Notwithstanding its non-overlapping activity spectrum, versikine ultimately derives from parental VCAN through ADAMTS- proteolysis (Fig. 1A): therefore, we hypothesized that VCAN expression (the substrate for versikine) and Batf3-DC abundance correlate in human cancer. To test this hypothesis, we carried out analysis of TCGA expression datasets across human carcinomas. We compared VCAN gene expression and Batf3-DC signature scores (Spranger et al., 2017) across 7591 samples from 20 TCGA cancer types (Fig. 1C and 1D). A significantly positive correlation between VCAN expression and Batf3-DC signature scores was observed in several human carcinomas (Fig. 1E), suggesting that the VCAN pathway regulates Batf3-DC density in multiple cancer types.

To help dissect mechanisms through which the VCAN pathway regulates DC, we generated novel *Vcan*-targeted models through genomic editing of *Vcan* N-terminal sequences (*Vcan* exons 3-6). The widely-used *Vcan*^hdf^ null mutant (hdf= heart defect) arose as the result of insertional mutagenesis into a site located 3’ to exon 7 encoding for a glycosaminoglycan-binding domain; this allele is embryonic lethal when reduced to homozygosity (Mjaatvedt et al., 1998). However, it is possible that residual versikine can be generated through partial expression of non-targeted, N-terminal *Vcan* sequences. To completely eliminate the possibility of versikine generation, we used CRISPR-Cas9-mediated mutagenesis to disrupt exon 3 sequences, thus abolishing transcription of all Vcan isoforms (and consequently, generation of versikine) (Fig. 1F). We derived two founders (Vcan1053, Vcan1058) bearing *Vcan* exon 3 deletions (16bp and 47bp, respectively) as shown in Supp. Fig. S1A/B. We validated a functional defect in *Vcan* message induction through stimulation of bone marrow-derived macrophages (BMDM) with lipopolysaccharide (LPS) (Supp. Fig. S1C). BMDM stimulated with LPS have been shown to preferentially transcribe V1 isoform, the precursor to versikine (Chang et al., 2014). The Vcan1053 transgenic line demonstrated the more severe defect in *Vcan* message stability and this line was subsequently selected for experiments in this study (hereafter designated *Vcan*+/-).

Lewis Lung Carcinoma (LLC) cells produce Vcan cell-autonomously (Kim et al., 2009). We knocked down endogenous Vcan expression in LLC cells using shRNA targeting *Vcan* exon 8 (encoding for the GAG*β* domain in versikine’s precursor V1 isoform, the major isoform produced in LLC - Fig. 1A), hereafter referred to as LLC^VcanKD^. We validated reduced transcription of Vcan in LLC^VcanKD^ cells using both 5’ and 3’ *Vcan* primers, as demonstrated in Supp. Fig. S1D. We characterized the intratumoral immune contexture in LLC^VcanKD^ tumors implanted in *Vcan*+/- mice through mass cytometry and compared to WT controls (Fig. 1G and Supp. Table S2). Vcan depletion resulted in expansion of CD8+ cells, consistent with the known role of non-proteolyzed VCAN in T-cell exclusion (Evanko et al., 2012; Gorter et al., 2010; McMahon et al., 2016).

Consistent with our primary hypothesis, we observed Batf3-DC loss in Vcan-depleted tumors. To refine the mass cytometry findings, we delineated intratumoral DC through 9-color flow cytometry as described by the Van Ginderachter group (Laoui et al., 2016) (Supp. Fig. S1E for gating strategy). Using this strategy, Batf3-DC in the cDC gate were depleted in LLC^VcanKD^ tumors implanted in *Vcan*+/- mice (Fig. 1H/I). Ectopic expression of versikine in LLC^VcanKD^ cells restored intratumoral Batf3-DC abundance to physiological levels (Fig. 1H/I). Intratumoral DC absolute count ratios corroborated the cell frequency findings: despite fluctuations in total cDC abundance across genotypes and individual tumors (Supp. Fig. S1F), Batf3-DC were selectively lost when Vcan was depleted in the tumor microenvironment (Fig. 1H/I). These results demonstrate that the stromal matrikine, versikine, is necessary and sufficient for Batf3-DC abundance.

### Stromal remodeling products promote an immunogenic TME *in vivo*

VCAN proteolysis in human cancers is a composite event that produces two simultaneous, coupled consequences: firstly, reduction in levels of parental VCAN and secondly, the novel activities of the derived matrikine, versikine. To uncouple versikine’s activity from the effects of parental VCAN depletion, we generated LLC cells stably expressing hemagglutinin (HA)-tagged versikine in the WT background (Fig. 2A). Expression of versikine in LLC cells did not result in grossly visible increase in angiogenesis or hemorrhagic propensity (Fig. 2B). Using an antibody against the HA tag, we determined that ectopically-expressed versikine was readily detectable by western blotting in murine tumor lysates at the expected MW of approximately 75kD (Fig. 2C).

**Fig. 2.**
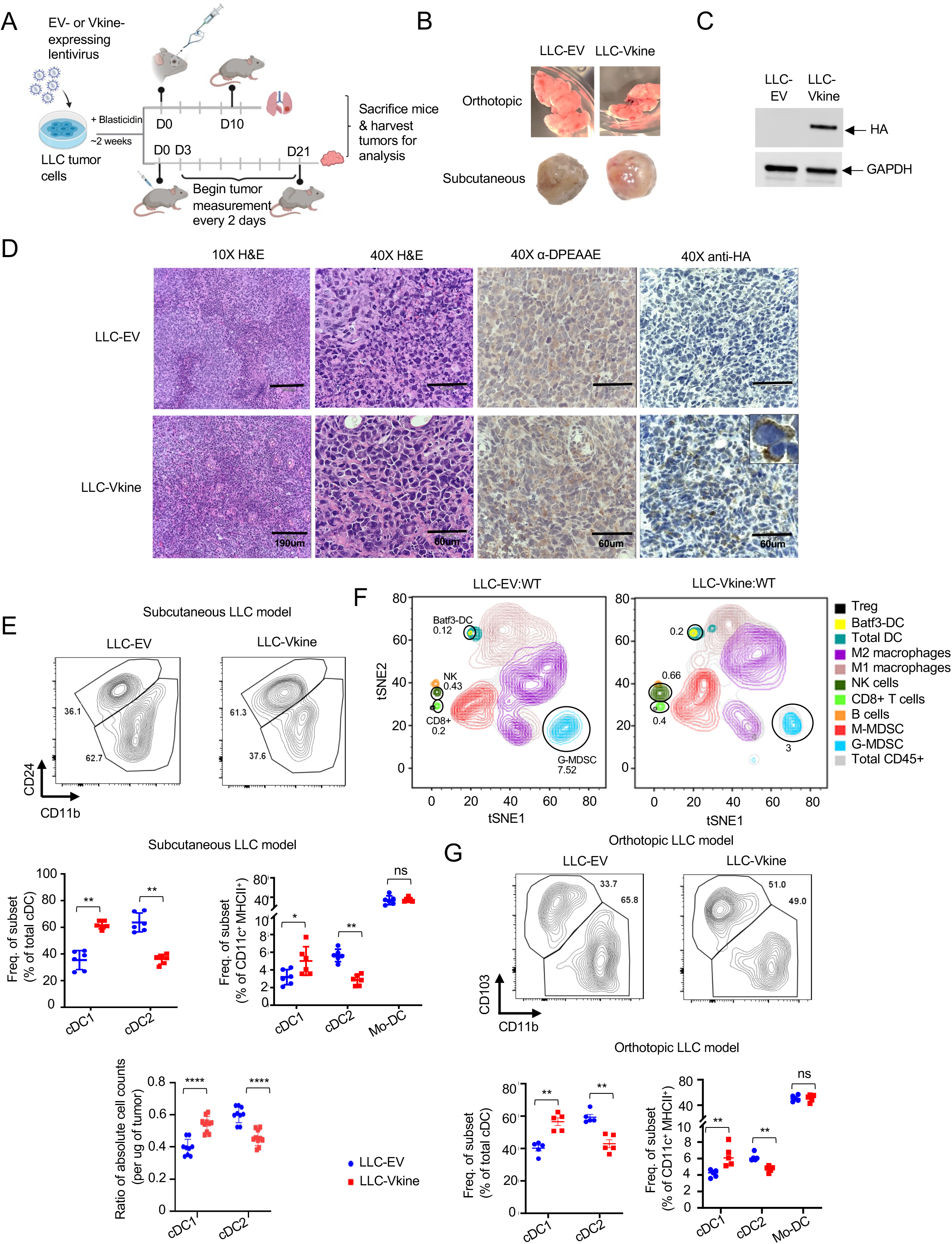
Stromal remodeling products promote an immunogenic TME *in vivo*. A: Experimental layout. LLC tumor cells were engineered to express HA-tagged versikine (LLC-Vkine) and empty vector controls (LLC-EV). 5x10^5^ LLC-Vkine or -EV cells were injected either subcutaneously (s.c.) on the flank or intravenously using retro-orbital approach. Tumor growth was measured every 2 days for subcutaneously injected LLC mice. After 21 days mice were sacrificed and tumors harvested for further analysis. For the orthotopic LLC model, mice were sacrificed after 10 days of tumor cell injection and lungs were harvested for further analysis. B: Gross morphology of orthotopic (top) and subcutaneous (bottom) LLC-EV and LLC-Vkine tumors. C: Anti-HA western blotting detects a 75kD band in LLC-Vkine tumor lysates, consistent with versikine. D: Representative immunohistochemistry (IHC) images showing α-DPEAAE and HA-tag staining of LLC-EV and LLC-Vkine tumors. Endogenous DPEAAE proteolysis is low-level and similar between LLC-EV and LLC-Vkine. Anti-HA staining localizes in a membranous distribution in LLC- Vkine cells (inset, larger magnification). E: Flow cytometric analysis of cDC subsets in subcutaneous LLC-EV and LLC-Vkine tumors. Gating strategy per Van Ginderachter group (Laoui et al., 2016), as delineated in Supplementary Fig. 1E. Quantification of cDC and TADC frequency (top of panel) and absolute count ratios (cDC1/cDC1+cDC2 and cDC2/cDC1+cDC2) (bottom of panel). F: Comparison of immune contexture (CD45+ cells) in LLC-EV vs. LLC-Vkine tumors by 31-marker mass cytometry. G: Flow cytometric analysis of cDC subsets in orthotopic LLC-EV and LLC-Vkine tumors (lung metastases induced by intravenous injection). Summary of cDC and TADC subset frequencies depicted to the right. Data are represented as mean±SD and are from one of three independent experiments with n=5 or 6 for each group. *p<0.05; **p<0.01;***p<0.001.

Murine transplantable tumors do not recapitulate the architecture of epithelial nests and stromal sheets; this limitation has been previously attributed to the acquisition of mesenchymal attributes by the tumor cells (Guerin et al., 2020) (Fig. 2D). However, the LLC model does retain physiological relevance due to its tumor-intrinsic production of Vcan that regulates myeloid cells in the TME (Kim et al., 2009). In this regard, LLC models a subset of human lung cancers with detectable VCAN production and VCAN processing in both stromal and epithelial compartments (Supp. Fig. S2B; negative controls in Supp. Fig. S2C). Ectopically-expressed versikine was detected in a membranous distribution consistent with its accumulation in the pericellular glycocalyx (Fig. 2D), a physiological site of VCAN cleavage (Hattori et al., 2011). Similar distribution of ectopic versikine was seen in B16 melanoma tumor cells that transcribe very low to undetectable endogenous *Vcan* message (Supp. Fig. S2A).

There were no differences in growth rates between LLC-EV and LLC-Vkine tumors (Supp. Fig. S2D). We analyzed cDC populations by conventional flow cytometry (Supp. Fig. S1E). Within the cDC gate, we confirmed Batf3-DC expansion in LLC-Vkine tumors (Fig. 2E), notably the opposite phenotype to that of Vcan depletion (Fig. 1H). There was no change in Mo-DC frequency between LLC-Vkine and LLC-EV tumors (Fig. 2E). By mass cytometry, in addition to Batf3-DC accumulation, we observed the expansion of an innate lymphoid NK1.1+NKp46+ population, increase in intratumoral CD8+ T cells as well as G-MDSC depletion (Fig. 2F). We previously hypothesized an impact of versikine on G-MDSC (Hope et al., 2016), based on known regulation of this population by versikine’s target, IRF8 (Waight et al., 2013). To confirm that Batf3-DC expansion was not peculiar to the non-orthotopic microenvironment of subcutaneous LLC, we injected EV- and versikine-expressing LLC cells intravenously and harvested lungs bearing metastatic deposits at D10 post-analysis, prior to clinical demise (Fig. 2G). We analyzed intratumoral DC through a similar flow cytometry strategy with the substitution of Batf3-DC marker CD103 for CD24: as per prior reports, we found CD103 to be more consistent in native lung tissue (Cabeza-Cabrerizo et al., 2021; Misharin et al., 2013). Similar to what we observed in subcutaneous LLC tumors, orthotopic lung LLC-Vkine tumors showed enhanced Batf3-DC (Fig. 2G).

To test whether versikine’s activities were limited to specific tumor types or murine genetic backgrounds, we extended our observations to other tumor models. Versikine promoted Batf3- DC in the orthotopic immunocompetent breast carcinoma model, 4T1, propagating in the Balb/c background (Supp. Fig. S2E). A preponderance of Batf3-DC in the cDC gate was clearly observed in 4T1-Vkine tumors. Growth rates of 4T1 versikine-replete tumors did not differ from those of their EV counterparts (Supp. Fig. S2F). We earlier reported a role for VCAN proteolysis in shaping the human bone marrow myeloma microenvironment (Hope et al., 2016). More recently, we developed the first Ras-driven immunocompetent myeloma model, VQ (Wen et al., 2021), a model that allows gene transfer into myeloma cells and engraftment into immunocompetent syngeneic recipients in the C57BL6/J background. Versikine-replete VQ myeloma tumors demonstrated enhanced Batf3-DC (Supp. Fig. S2G), whereas clinical progression was unaffected by versikine (Supp. Fig. S2H). Therefore, stromal matrikines can promote pro-immunogenic TME in both solid and hematopoietic cancers.

### Pre-DC differentiation is unaffected by stromal matrikine signaling

To explain how stromal remodeling products can support Batf3-DC, we first tested a hypothesis focused on differentiation of uncommitted tumor-seeding pre-DC (Diao et al., 2010). Indeed, our observation that recombinant versikine promoted the generation of Batf3-DC from mouse BM treated with Flt3L would seem to support this hypothesis (Hope et al., 2017). *Irf8* and other Batf3- DC “signature” transcripts (*Irf8*, *Batf3*, *Cxcl9*, *Cxcl10*) were increased in the bulk transcriptome of versikine-replete (LLC-Vkine) tumors (Fig. 3A). Irf8 is a “terminal selector” for the Batf3-DC lineage (Sichien et al., 2016). *Batf3*-transcript increase corroborates versikine-induced Batf3-DC abundance because *Batf3* expression range is very narrow (Supp. Fig. S3A and Supp. Table S3). *Id2* transcripts did not differ between versikine-replete and EV-tumors but *Id2* is more broadly expressed and not highly expressed in Batf3-DC (Supp. Fig. S3A and Supp. Table S3).

**Fig. 3.**
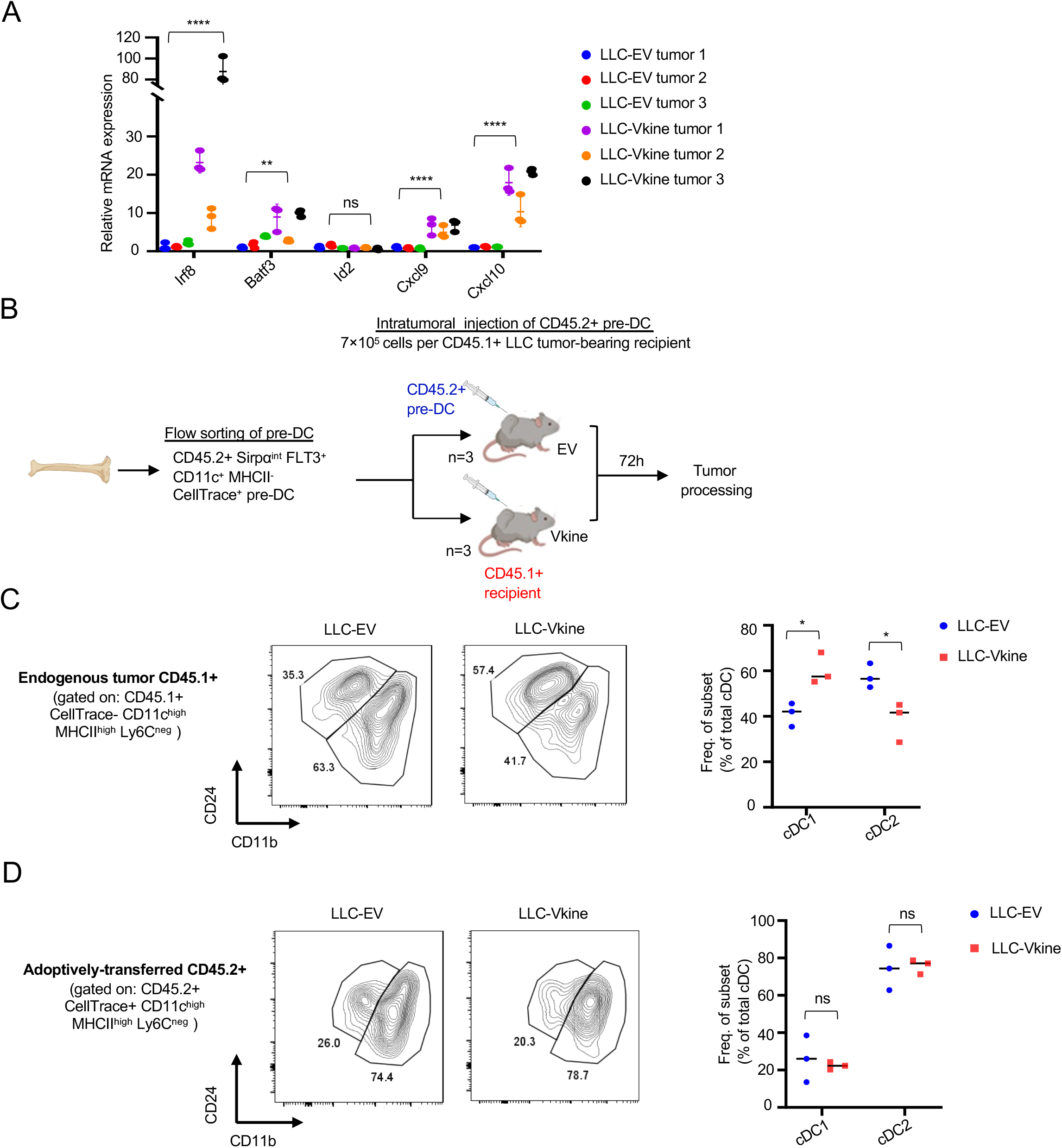
Pre-DC differentiation is unaffected by stromal signals. A: RT-PCR analysis for Batf3- DC “signature” genes in bulk LLC-EV and LLC-Vkine tumor mRNA. B: Schematic layout of the pre-DC adoptive transfer experiment. Pre-DC were harvested from BM of Flt3L-mobilized CD45.2+ mice and adoptively transferred into LLC-EV or LLC-Vkine tumors implanted in CD45.1+ recipients. 72 hours post adoptive transfer, tumors were harvested, processed and cDC subsets were analyzed by flow cytometry. C: Representative flow plots showing CD45.1+ endogenous cDC subset frequencies (left) and quantifications depicted to the right. D: Flow plots showing CD45.2+ adoptively-transferred cDC subset frequencies (left) and quantifications (right). In (A), (C) and (D) ns, non-significant, *p<0.05; **p<0.01;***p<0.001; ****p<0.0001.

To test our differentiation hypothesis, we sorted CD45.2+ pre-DC precursors from the BM of *in vivo* Flt3L-mobilized mice (Supp. Fig. S3B). CD45.2+ pre-DC were adoptively-transferred intratumorally into subcutaneous LLC-EV and LLC-Vkine tumors implanted in CD45.1+ recipients (Fig. 3B). At 72 hours post-adoptive transfer, tumors were dissociated and CD45.2+, as well as endogenous CD45.1+, DC fractions were enumerated and characterized by flow cytometry. The CD45.1+ endogenous cDC composition served as an internal control for the experiment. As expected, CD45.1+ endogenous Batf3-DC were increased in LLC-Vkine tumors (Fig. 3C). By contrast, CD45.2+ Batf3-DC and cDC2 did not differ between LLC-Vkine and -EV controls (Fig. 3D). Thus, stromal remodeling does not appear to act by forcing the differentiation choice of pre-DC (Batf3-DC vs. cDC2) that seed peripheral tumor sites, at least within the time-frame of our *in vivo* adoptive transfer differentiation assay.

### Distinctive DC activation program by the stromal matrikine, versikine

Our finding that versikine did not impact on short-term pre-DC differentiation raised alternative hypotheses, potentially implicating Batf3-DC survival or recruitment. Specifically, we questioned whether versikine activated survival mechanisms in Batf3-DC, either in cell-autonomous or non-cell-autonomous manner. Non-cell autonomous survival pathways could be mediated through innate lymphoid cells (NK or ILC1) or other supporting cell types in the TME. We earlier demonstrated an expansion of NKp46+NK1.1+ cells in versikine-replete tumors (Fig. 2F), suggesting that versikine might constitute an upstream activator of Batf3-DC cross-talk with supporting innate lymphoid cells.

To explore versikine-induced activation pathways in Batf3-DC, we took avail of Batf3-DC cell model, MutuDC1940 (Fuertes Marraco et al., 2012). These cells have been shown to constitute *bona fide* Batf3-DC equivalents, in terms of immunophenotype, transcriptional factor profile, cytokine secretion and cross-presentation capacity (Fuertes Marraco et al., 2012). To explore steady-state changes in MutuDC1940 transcriptome in the presence of versikine, we generated stable MutuDC1940-Vkine cell lines through lentiviral transduction (Fig. 4A). Stable expression of versikine did not alter the baseline growth characteristics of MutuDC1940 cells (data not shown). MutuDC1940-versikine cell morphology was not significantly different from MutuDC1940-EV counterparts, albeit with slightly more developed dendritic appearance (Fig. 4B).

**Fig. 4.**
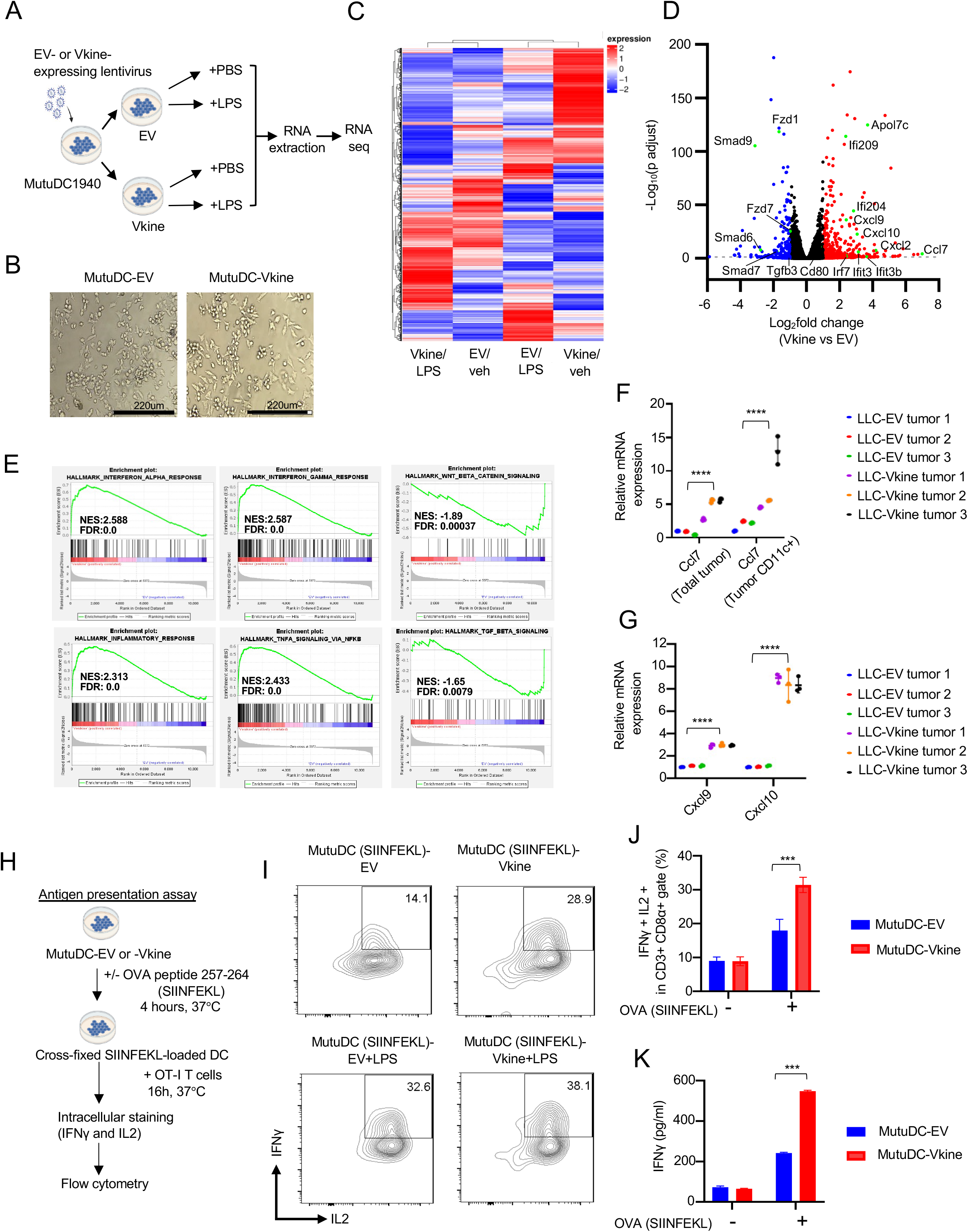
Distinctive DC activation by the stromal matrikine, versikine. A: Schematic layout of experiment. Stably-expressing MutuDC1940-EV or -Vkine cells were additionally stimulated for 4hs with vehicle (PBS) or LPS (100 ng/mL). At the end of stimulation, cells were lysed in RNA buffer for downstream RNA extraction and processing. B: Gross morphology of MutuDC1940 cells engineered to express versikine (Vkine) or empty vector (EV). Phase contrast, 10X magnification, scale bar 220µm. C: Hierarchical clustering of transcriptomic profiles by RNAseq analysis of MutuDC1940 cells transduced with EV or versikine (Vkine) and additionally stimulated with TLR4 agonist lipopolysaccharide (LPS) or vehicle (PBS). D: Volcano plot highlighting key differentially expressed genes in MutuDC1940-Vkine vs. -EV cells. E: Gene-Set Enrichment Analysis (GSEA) of significantly upregulated (left, middle) and downregulated (right) pathways in MutuDC1940- Vkine vs. -EV cells. F: RT-PCR analysis for *Ccl7* message in bulk LLC-EV and LLC-Vkine tumor mRNA (left) and CD11c+-selected (magnetically-separated) DC mRNA (right). G: RT-PCR analysis for *Cxcl9 and Cxcl10* message in RNA extracted from CD11c+-selected (magnetically-separated) cells derived from LLC-EV or LLC-Vkine tumors. H: Schematic layout of antigen-presentation experiment. I: Flow cytometry for endogenous IFN*γ* and IL2 of OT-I CD8+ T cells co-cultured with SIINFEKL-loaded MutuDC1940, either -EV or -Vkine, with or without LPS. J: Quantitation of OT-I flow cytometric analysis of antigen presentation assay. K: IFN*γ* by ELISA in supernatants from OT-I+ MutuDC1940:SIINFEKL co-cultures in the antigen presentation assay. ns, non-significant, *p<0.05; **p<0.01;***p<0.001; ****p<0.0001.

Versikine elicited a cell-autonomous co-stimulatory transcriptional program in MutuDC1940 cells distinct from the transcriptional program elicited by Toll-like receptor 4 (TLR4) agonist, lipopolysaccharide (LPS) (Fig. 4C). Combination of versikine+ LPS elicited a transcriptional signature distinct from either stimulus alone (Fig. 4C). We subsequently characterized the unique transcriptional output of versikine in MutuDC1940 cells. Examination of the genes and pathways triggered by versikine suggests a clear stimulatory role (Supp. Fig. S4A). Upregulated genes were involved in DC maturation (interferon-stimulated genes such as Ifi209 and Ifi204), chemokines (*Ccl7, Ccl2, Cxcl9, Cxcl10*) and co-stimulatory ligands (*Cd80, Cd40*)(Fig. 4D and Supp. Table S4). Downregulated genes include components of TGF*β* signaling and Wnt signaling, both associated with immunosuppression (Conejo-Garcia et al., 2016). Furthermore, Gene Set Enrichment Analysis (GSEA) demonstrated that versikine promotes transcriptional changes associated with immunogenic polarization of Batf3-DC, e.g. enrichment of IFN-α response, IFN-γ signaling, NF-*κ*B-induced TNF signaling, inflammation; and downregulation of immunosuppressive pathways such as Wnt-β-Catenin and TGFβ signaling (Fig. 4E). Several of the top hits were confirmed by RT-PCR and at the protein level, by ELISA (Supp. Fig. S4B/C/D/E). Moreover, hits upregulated at steady-state through stable introduction of versikine constructs were also confirmed following exposure of MutuDC1940 cells to supernatant isolated from cultures of HEK293 cells secreting recombinant versikine (Supp. Fig. S4F). One of the top-induced versikine-signature genes, Ccl7, was recently shown to act as a Batf3-DC chemoattractant (Zhang et al., 2020). Indeed, bulk LLC-Vkine tumors expressed higher Ccl7 message than LLC-EV tumors (Fig. 4F), as did immunomagnetically-separated CD11c+ cells from LLC-Vkine tumors (Fig. 4F).

We subsequently sought to directly interrogate the functional consequences of the activation program conferred by versikine alone or following co-stimulation with TLR4 agonist, LPS. Versikine upregulated co-stimulatory receptors (B7 receptors, CD40) as well as co-stimulatory cytokines in MutuDC1940 either alone or in association with LPS (Fig. 4D). These results raised the possibility that versikine, alone or in combination with other maturation signals, promotes antigen-presenting capacity. To test this hypothesis, we carried out antigen-presentation assays using the OVA (ovalbumin) antigen system in conjunction with T-cell receptor-engineered OT-I T cells (Fig. 4H). Versikine- and EV- MutuDC1940 cells were pulsed with SIINFEKL peptides and co-cultured with OT-I cells. The results are shown in Fig. 4I/J/K and Supp. Fig. 4G/H. Versikine alone more than doubled the percentage of primed OT-I cells secreting IFN*γ* and IL-2 by flow cytometry, confirmed through ELISA of the culture supernatant for IFN*γ*. Providing both versikine and LPS maximized T-cell priming. These results further demonstrated that stromal remodeling products can synergize with immunogenic danger signals to maximize stimulatory DC antigen presentation and T-cell priming.

### Batf3-DC accumulation requires atypical innate lymphoid support

The results in the previous section demonstrated that stromal matrikines can activate Batf3-DC cell-autonomously. To determine whether this activation program regulates Batf3-DC/ NK cross- talk *in vivo*, we explanted CD11c+ DC from primary versikine-replete vs. EV- tumors through immunomagnetic separation. Freshly explanted CD11c+ cells from LLC-Vkine tumors expressed higher levels of NK-regulating IL-23 (*α* subunit), IL-27 (p28 and EBI3 subunits) and IL-15 (Fig. 5A). These results demonstrated that the distinct versikine-induced Batf3-DC activation program incorporated an NK-activating module.

**Fig. 5.**
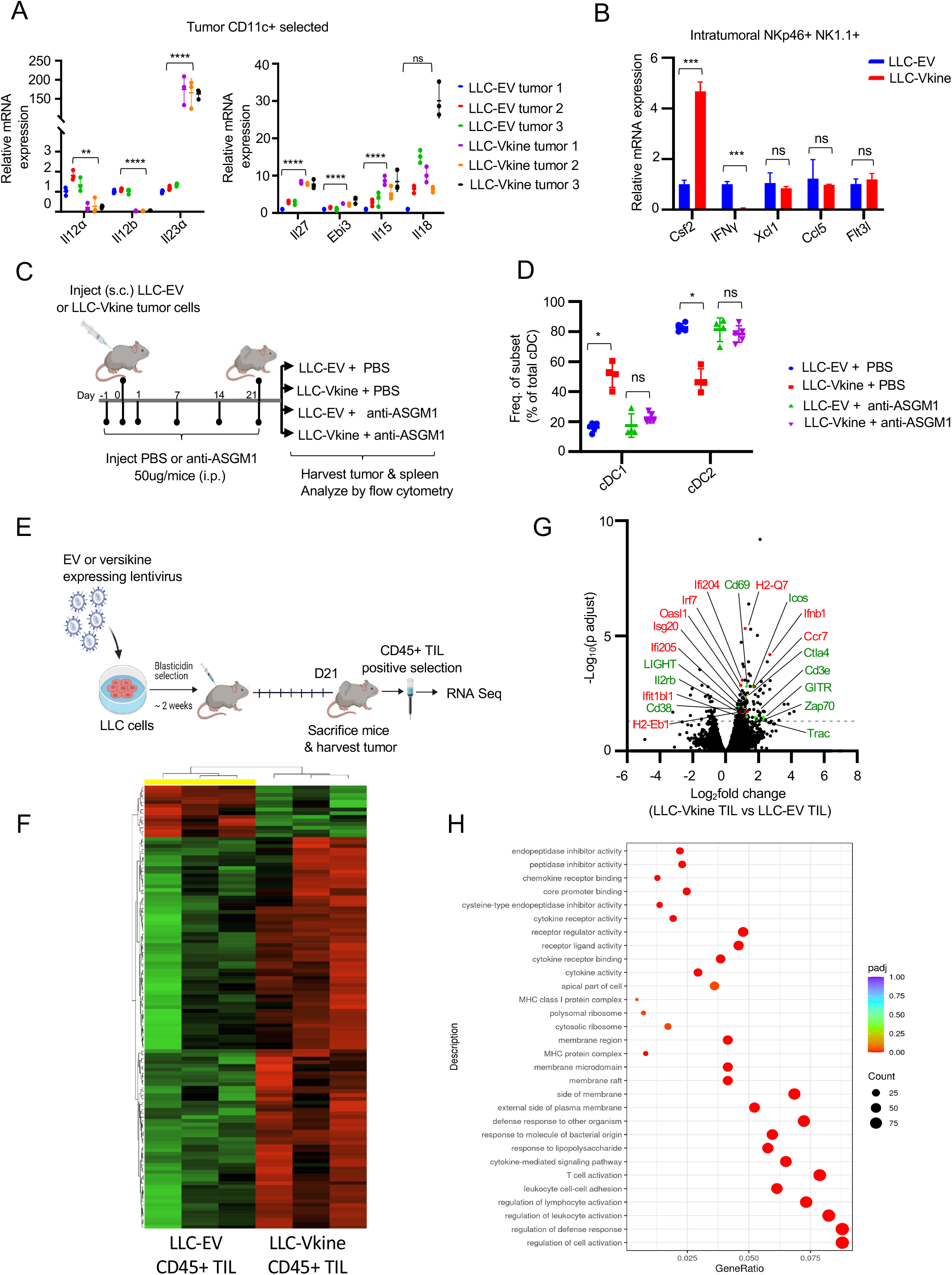
Batf3-DC accumulation requires atypical innate lymphoid support. A: RNA from *ex vivo*-magnetically separated CD11c+ cells from LLC-EV and LLC-Vkine tumors was subjected to RT-PCR to measure relative expression of NK-activating cytokines (as shown). Data are represented as mean±SD, n=3 for each group, B: RNA from flow-sorted NKp46+NK1.1+ cells flow-sorted from LLC-EV and LLC-Vkine tumors was subjected to RT-PCR using gene primers, as shown. C: Schematic layout of NK-depletion. D: Summary of cDC subset frequency by flow cytometric analysis in LLC-EV vs. LLC-Vkine tumors following treatment with NK-depleting antibody (anti-ASGM1) or vehicle (PBS). E: Layout of experiment to characterize immune infiltrates in LLC-EV vs. LLC-Vkine tumors. F: Hierarchical clustering of transcriptomic profiles by RNAseq analysis of CD45+ tumor-infiltrating leukocytes (TIL) extracted from LLC-EV vs. LLC- Vkine tumors. G: Volcano plot highlighting key differentially expressed genes in CD45+ TIL from LLC-Vkine tumors compared to LLC-EV tumors. Genes whose overexpression has been linked with APC activation in red, genes whose overexpression has been linked with T cell activation in green. H: Gene Ontology (GO) analysis of pathways enriched in CD45+ cells from LLC-Vkine vs. LLC-EV tumors. Data are represented as mean±SD, ns, non-significant, *p<0.05; **p<0.01;***p<0.001; ****p<0.0001.

Given the fact that versikine induced an NK- activating module (Fig. 5A) and resulted in expansion of a NKp46+NK1.1+ innate lymphoid subset (Fig. 2F), we sought to further characterize the innate-lymphoid program induced by versikine. Previous studies implicated NK-derived differentiation/survival mediator Flt3L as well as chemo-attractants Xcl1 and Ccl5 in Batf3-DC support (Barry et al., 2018; Bottcher et al., 2018). Moreover, NK-derived IFN*γ* was recently shown to induce IRF8, a Batf3-DC “terminal selector” (Lopez-Yglesias et al., 2021). To determine whether any of these previously reported mechanisms were relevant, we flow-sorted NKp46+NK1.1+ cells from LLC-EV and LLC-Vkine tumors. Surprisingly, NKp46+NK1.1+ cells from LLC-Vkine tumors were potent expressors of GM-CSF and relatively weak expressors of IFN*γ* compared to LLC-EV-derived cells (Fig. 5B). Expression of *Xcl1*, *Flt3l* and *Ccl5* remained unchanged (Fig. 5B). Therefore, versikine results in expansion of an atypical NK subset expressing low IFN*γ* despite high cytotoxicity receptor (NKp46) expression, and robustly expressing GM-CSF, an essential survival factor for Batf3-DC both at steady-state and in the tumor microenvironment (Greter et al., 2012). These results suggest that versikine engages innate lymphoid cells through a previously unreported mechanism in the TME. These cells are, however, reminiscent of a very recently reported spleen-resident ILC1-like subset that nurses Batf3-DC and promotes CD8+ priming in viral infection (Flommersfeld et al., 2021).

We further sought to determine whether versikine requires innate lymphoid cells to promote Batf3- DC. We used asialo-GM1 antibody for *in vivo* depletion because we hypothesized that the atypical innate lymphoid cell subset induced by versikine is more closely related to circulating NK cells seeding the tumor, a hypothesis consistent with the findings in (Flommersfeld et al., 2021). The layout of the experiment is presented in Fig. 5C. Asialo-GM1+ depletion was approximately 70% efficient (Supp. Fig. S5A) and completely abrogated versikine-mediated enhancement of Batf3- DC (Fig. 5D and Supp. Fig. S5B).

### TLR2 and CD44 are dispensable for Batf3-DC accumulation

Versikine’s receptor is unknown but two major contenders merited direct interrogation: TLR2 and CD44. Non-proteolyzed VCAN is thought to act through TLR2 (Kim et al., 2009; Tang et al., 2015) but it is unclear if versikine utilizes TLR2. However, *Tlr2* loss had no impact on versikine-induced Batf3-DC enhancement (Supp. Fig. S5C/D/E).

VCAN’s N-terminal link domains bind hyaluronan (Fig. 1A). In previous work from our group, we demonstrated bioactivity of recombinant versikine on myeloid cells in the absence of bound hyaluronan (Hope et al., 2016). However, it remained formally possible that versikine’s effects are mediated through hyaluronan receptors *in vivo*, chief among which is CD44 (Misra et al., 2015). Moreover, versikine could bind directly to CD44, through its link domains. Stromal *Cd44* loss made no difference to versikine-induced Batf3-DC enhancement (Supp. Fig. S5F/G/H).

### Robust T-cell activation in response to stromal signals *in vivo*

To further understand mechanisms by which stromal remodeling products rewire tumor-infiltrating immune cell networks *in vivo,* we compared the transcriptomic profiles of CD45+ tumor-infiltrating leukocytes (TIL) isolated from versikine-replete vs. control LLC tumors (Fig. 5E).

Versikine radically remodeled the immune microenvironment in LLC tumors (Fig. 5F and Supp. Table S5). Immune infiltrates in versikine-replete microenvironments expressed hallmarks of APC activation (upregulation of *MHCII, Ccr7, Ifnb1, Irf7* and several interferon-responsive genes) and a compelling T-cell co-stimulation and activation signature (*Cd69, Ctla-4, Icos, Zap70, Il2rb, Cd38, Light, Gitr*). Moreover, there was a significant increase in T-cell-specific transcripts (*CD3e*, TCR genes), consistent with CD8+ T cell expansion seen by mass cytometry (Fig. 5G). The upregulation of TNF family member ligand Light (Tnfsf14) in particular, is noteworthy given the association between this co-stimulatory pathway and tertiary lymphoid structure formation (Johansson-Percival et al., 2017), where peritumoral Batf3-DC have been previously reported in some cases (Lavin et al., 2017). The results further demonstrate that in addition to the quantitative changes shown by mass cytometry in Fig. 2F, versikine promotes a robust T-cell activation program.

### Stroma-licensed Batf3-DC are “poised” and hypersensitive to nucleic acid sensing *in vivo*

Previous studies have highlighted the paradox of accumulation of immunogenic DC along the tumor rim (Pai et al., 2020), but no compelling mechanism has emerged. We hypothesized that stroma-licensed DC become “poised” to respond to physiological, endogenous maturation signals arising from necrotic cells. The cGAS/STING pathway, a sensor of exogenous double-stranded DNA, has emerged as a central mediator of innate sensing of tumors (Corrales et al., 2015). We reasoned that versikine-activated DC may respond to limiting doses of STING agonist (Fig. 6A). LLC-EV and LLC-Vkine tumors were challenged with *sub*-therapeutic doses of STING agonist DMXAA (150-200 mcg) or vehicle (NaHCO3). Notably, Gajewski and colleagues established dose-response relationships for intratumoral DMXAA- the MTD was 500 mcg, as unacceptable toxicity was observed at higher doses (Corrales et al., 2015). Tumor response curves are shown in Fig. 6B and survival plots (Kaplan-Meier) in Fig. 6C. EV-tumors did not appreciably respond to vehicle or subtherapeutic STING agonist (DMXAA200). By contrast, versikine lowered the response threshold to DMXAA so that versikine-replete tumors demonstrated a consistent response to single doses of *sub*therapeutic DMXAA. Several versikine-replete tumors developed necrotic eschars by 24 hours after subtherapeutic DMXAA injection (Fig. 6D). By contrast, none of the control mice developed an eschar within that time frame or with similar consistency. To determine whether versikine reduced therapeutic threshold through a classical type I-IFN response to DMXAA, we harvested tumors for RNA extraction at 2 hours post-DMXAA. Versikine- replete tumors demonstrated several-fold increase in interferon-*α* transcripts (particularly IFN*α*2 and IFN*α*4), and to a lesser degree IFN*β*1 transcripts (Fig. 6E). A fuller list of up- and down- regulated genes in this experiment is provided in Supp. Table S6. These results demonstrate that versikine lowered the threshold for a classical, type-I interferon-mediated, STING agonist response.

**Fig. 6.**
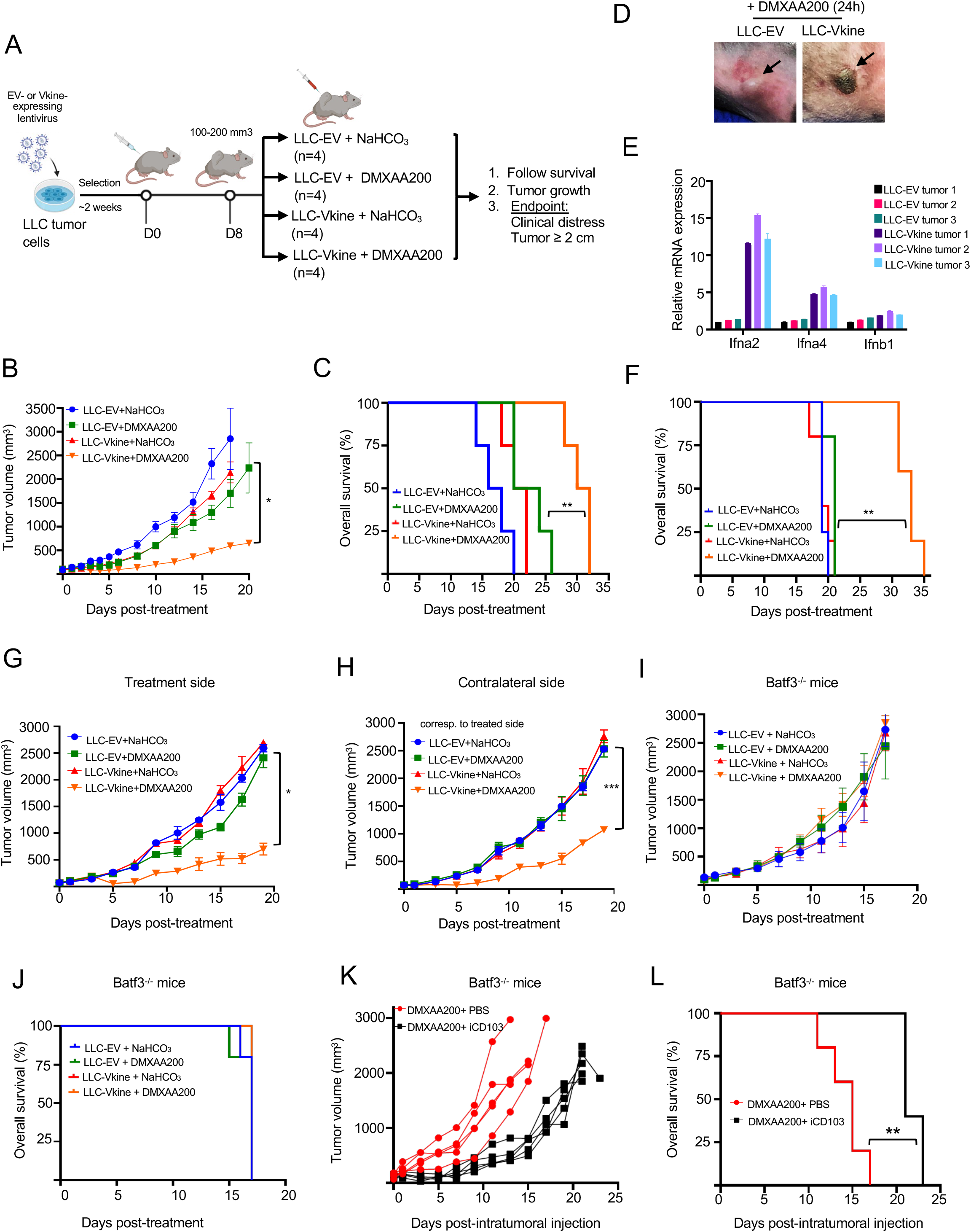
Stroma-licensed Batf3-DC are “poised” and hypersensitive to nucleic acid sensing *in vivo*. A: Schematic layout of the experiment. 5×10^5^ LLC-EV or LLC-Vkine cells were injected (s.c.) in the mouse flank and on day 8 (tumor volume 100-200mm^3^) tumors received a single intratumoral (IT) injection of either vehicle (NaHCO3) or sub-therapeutic dose (200 mcg) of STING agonist, DMXAA. Tumor growth was monitored every 2 days and mice were sacrificed when they showed clinical distress or tumor burden > 2cm^3^. B: Growth curves of LLC-EV and LLC-Vkine tumors challenged with a single subtherapeutic dose (200 mcg) of IT DMXAA on Day 0 (DMXAA200) or vehicle (NaHCO3). C: Kaplan-Meier survival curves for the experiment in (B), **=p<0.01 by log-rank test. D: Representative images showing development of hemorrhagic necrosis and a necrotic eschar in LLC-Vkine, but not LLC-EV tumors, at 24 hours post-IT DMXAA200. E: Transcriptomic analysis of LLC-EV and LLC-Vkine tumors harvested at 2 hours post-IT DMXAA200. See also Supp. Table. S6. F: Versikine-DMXAA synergy generates an abscopal effect in LLC tumors that produces a survival advantage. **=p<0.01 by log-rank test. G: Growth curves of <u>treatment-side</u> LLC-EV and LLC-Vkine tumors challenged with a single subtherapeutic dose (200 mcg) of IT DMXAA on Day 0 (DMXAA200) or vehicle (NaHCO3). H: Growth curves of <u>contralateral side</u> unmanipulated LLC tumors, according to corresponding treatment side configuration (treatment as in Panel G). I: Response to DMXAA200 is lost in *Batf3*-/- recipients. Growth curves of LLC-EV and LLC-Vkine tumors challenged with a single subtherapeutic dose (200 mcg) of IT DMXAA on Day 0 (DMXAA200) or vehicle (NaHCO3) in *Batf3*-/- recipients. J: *Batf3*-loss abrogates the survival advantage seen in WT type (panel 6C). K: Efficacy of DMXAA200 in LLC-Vkine tumors implanted in *Batf3*-/- recipients is restored following adoptive transfer of iCD103 (see also Supp. Fig. S6D). L: Adoptive transfer of iCD103 in LLC- Vkine tumors implanted in *Batf3*-/- recipients restores survival advantage of mice treated with DMXAA200. **=p<0.01 by log-rank test.

To determine whether low doses of STING agonist could still generate an abscopal effect, we studied mice bearing tumors inoculated in both flanks as delineated in Supp. Fig. S6D. The treated side was inoculated with EV or versikine-expressing LLC cells; the contralateral, non-treated, side was inoculated with unmanipulated LLC cells. Notably, ectopic versikine is bound in the pericellular halo (glycocalyx) (Fig. 2D) and likely does not circulate to an appreciable degree. We observed a consistent abscopal effect when versikine-replete tumors were injected with 200 mcg DMXAA (Fig. 6F/G/H). EV- tumors treated with the same subtherapeutic dose failed to elicit any response, either on the treatment or contralateral sides. STING agonist hypersensitivity produced consistent primary tumor and abscopal effects across genetic backgrounds, as we observed tumor responses and survival benefit in the breast carcinoma 4T1 orthotopic model (Supp. Fig. S6A/B/C).

Our hypothesis that stromal matrikines render Batf3-DC hypersensitive to nucleic acid sensing *in vivo* would predict that DMXAA200 would be ineffective in the absence of Batf3-DC. Thus, we repeated the experiment (as delineated in Fig. 6A) in *Batf3*-null recipients. As shown in Fig. 6I, DMXA200 was globally ineffective in the *Batf3*-null background and survival benefit was lost (Fig. 6J). To confirm that the responsible actors were Batf3-DC (rather than another Batf3-expressing lineage), we attempted to rescue the null phenotype with intratumoral adoptive transfer of iCD103, BM-derived primary Batf3-DC-equivalents generated in culture using the protocol by Merad, Sparwasser and collaborators (Mayer et al., 2014)(Supp. Fig. S6D). We confirmed iCD103 to have a consistent Batf3-DC-like phenotype in our hands (Supp. Fig. S6E). Adoptive transfer of iCD103 restored low-dose STING agonist efficacy (Fig. 6K). iCD103 adoptive transfer restored a survival benefit (Fig. 6L). To confirm the findings in a different C57BL6/J model, we chose B16 melanoma (the 4T1 model could not be used as *Batf3*-null animals were maintained in the C57BL6/J background). B16 tumors responded to subtherapeutic doses of DMXAA in the presence of versikine but not EV (Supp. Fig. S6F/G). Efficacy was lost in the *Batf3*-null background (Supp. Fig. S6H) but response to low-dose STING agonists was restored, at least in a subset of mice, when iCD103 were adoptively transferred (Supp. Fig. S6I). Notably, we speculate that engraftment of iCD103 was less consistent in B16 (compared to LLC) due to the immune infiltrate paucity in B16 (B16 are relatively “cold” tumors and are likely relatively refractory to engraftment of transferred DC). Our results demonstrate that stroma-licensed Batf3-DC are “poised” and hypersensitive to exogenous nucleic acid sensing.

### Stromal remodeling signals promote antigen-specific CD8+ responses *in vivo*

We then asked whether hypersensitivity of stroma-licensed DC to DNA sensing translates into enhanced antigen-specific effector responses *in vivo*. To this end, we employed the ovalbumin (OVA) system as an *in vivo* model antigen (Fig. 7A). EV- and versikine-expressing LLC cells were engineered to express full-length OVA (LLC-OVA). EV- or versikine-replete LLC-OVA tumors were challenged with therapeutic dose of DMXAA (500 mcg). Five days after challenge, spleens were harvested and analyzed by flow cytometry for antigen-specific effector responses using an antigen-specific tetramer assay. Challenge of versikine-replete tumors more than doubled the magnitude of the antigen-specific response in the CD8+ compartment as determined by MHCI:SIINFEKL-tetramer staining (Fig. 7B). Moreover, spleens contained a larger proportion of CD8+CD62L+CD44+ cells with a central memory phenotype (Fig. 7C). The results demonstrate that harnessing of stromal signals results in robust antigen-specific CD8+ effector responses *in vivo*.

**Fig. 7.**
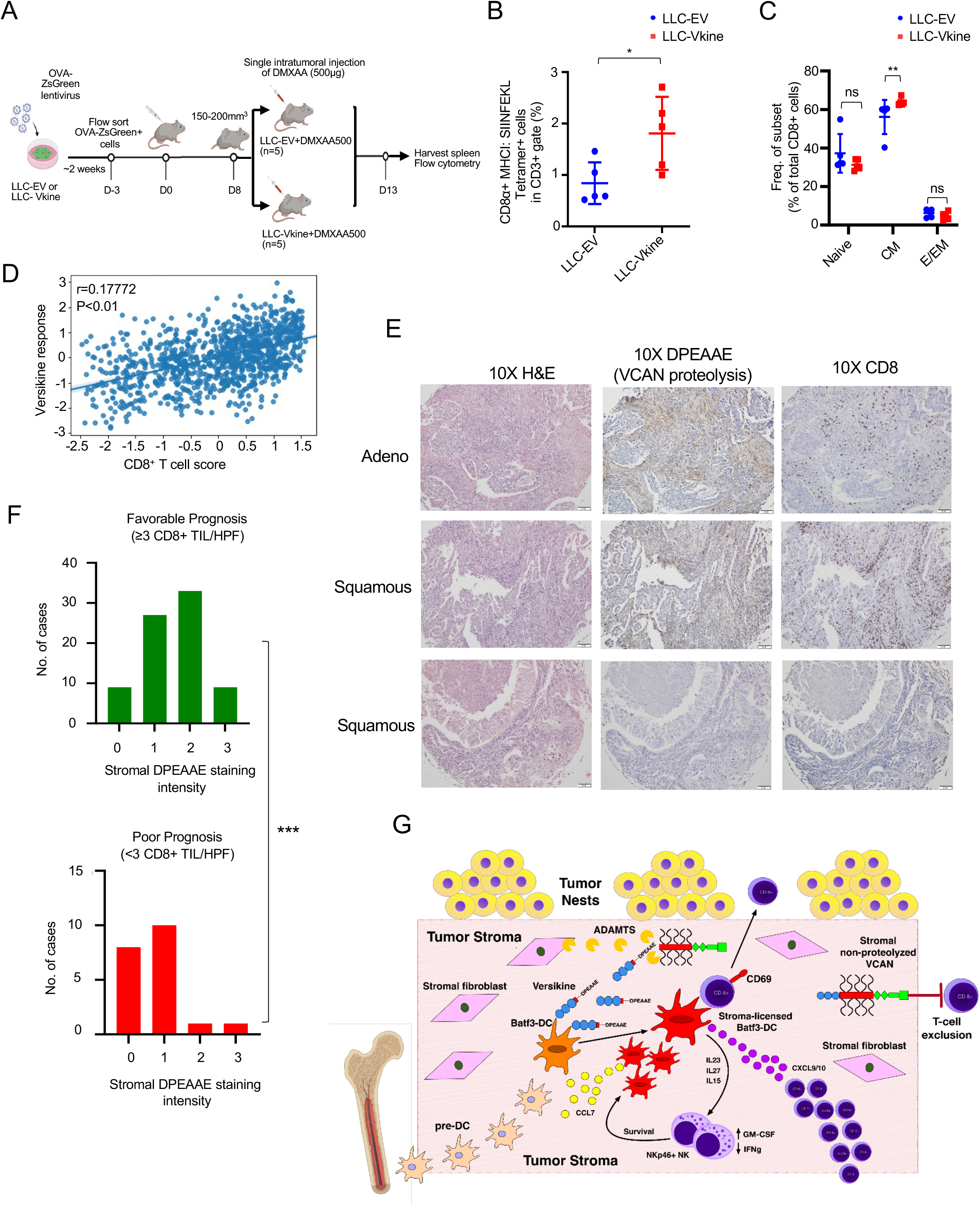
Stroma remodeling promotes antigen-specific CD8+ responses *in vivo* and correlates with CD8+ T cell infiltration in human lung cancer. A: Schematic layout of the *in vivo* antigen-specific response experiment. B: Frequency of MHCI:SIINFEKL tetramer+ CD8+ splenocytes in mice bearing LLC-EV vs. LLC-Vkine tumors at 5 days post-challenge with therapeutic dose of STING agonist (DMXAA500). C: CD8+ subset frequency in the spleen of mice treated as in panel 7B. Naïve (CD44-CD62L+), central memory [CM, (CD44+CD62L+)], effector/ effector memory [E/EM, (CD44+CD62L-)]. D: Correlation between *in vitro* versikine signature and CD8+ T cell scores. Significance measured using a linear model while accounting for total immune infiltration. E: DPEAAE staining in human lung cancers and associated CD8+ infiltration. DPEAAE is a neoepitope revealed through VCAN-V1 proteolysis at the site that generates versikine. F: Distribution of DPEAAE stromal staining intensity across lung cancer prognostic subgroups [pauci-immune (poor prognosis) and immune-rich (favorable prognosis) at cutoff 3 CD8+ TIL/HPF, p<0.001 by two-tailed Mann-Whitney test]. G: Schematic summary of stromal remodeling pathways regulating SDC abundance and function. Stroma-licensed Batf3-DC are supported by an atypical innate lymphoid subset expressing cytotoxicity receptors, low IFN*γ* and high GM-CSF. In our model, repriming of T-cell effectors by stromal-licensed DC results in T-cell proliferation, tumor infiltration and an “inflamed” tumor bed. Conversely, tolerogenic polarization of DC by non-proteolyzed VCAN (e.g., through TLR2) results in T-cell exclusion and stalling of effector T- cells at the tumor border.

### Stroma-licensed DC signature expression correlates with CD8+ scores in human lung cancer

Evidence linking stromal DC licensing with CD8+ density in human cancers has been lacking. To address this question, we generated a unique DC response signature from 200 genes whose expression was significantly altered in versikine-activated MutuDC1940 cells (Fig. 4C/D). We then correlated this stroma-licensed DC signature to CD8+ T-cell scores estimated in TCGA expression data for 1017 lung cancers (see Materials and Methods for details). The results are shown in Fig. 7D and Supp. Fig. S7A. We observed a significant, albeit weak, correlation between stroma-licensed DC transcriptional output and CD8+ scores. Despite the inherent limitations of the analysis (application of an *in vitro* generated signature using cultured mouse DC to primary bulk human tumor data), the results support a link between stromal DC activation and CD8+ density in human cancer.

### Stromal VCAN proteolysis correlates with CD8+ infiltration in human lung cancer

CD8+ infiltration has prognostic significance in human lung cancer (Zeng et al., 2016) as well as predictive significance for efficacy of checkpoint inhibitor-based immunotherapy (Fumet et al., 2018). A cutoff of 3-5 CD8+ cells/HPF has been used in some studies to designate CD8+TIL-rich vs. -poor tumors bearing favorable and unfavorable prognosis, respectively (see individual studies referenced in a meta-analysis (Zeng et al., 2016)). To determine whether stromal VCAN proteolysis may be associated with prognostic immune infiltration groups in human patients, we subdivided 98 NSCLC biopsies in our TMA into pauci-immune (0-2 CD8+ TIL/HPF in both stromal and epithelial compartments, n=26) and immune-rich (equal or greater than 3 CD8+ TIL/HPF in either stromal or epithelial compartments, n=72). The distribution of stromal DPEAAE staining intensity (0, 1, 2, 3 assessed by pathologist KAM) was compared between the groups. The results are shown in Fig. 7 E/F, Supp. Fig. S7B and Supp. Table S1. We found a statistically significant association between stromal VCAN proteolysis and CD8+ infiltration in human NSCLC.

## DISCUSSION

Our data address several unsettled conundrums in tumor DC biology: First, the persistence of a rare subset of stimulatory DC within the overwhelmingly suppressive microenvironments of established tumors, characterized by a vast excess of regulatory myeloid cells. Second, the enigmatic topography of SDC activity along tumor margins, distal to tumor nests. Third, the mechanisms regulating T-cell infiltration versus the T cell-excluded phenotype characterized by stalling of immune effectors within the peritumoral stroma. Fourth, the clinical impetus to increase Batf3-DC density within tumors and promote infiltration by natural and engineered T-cell and other immune effectors.

Batf3-DC displacement from tumor nests has been associated with activating tumor-intrinsic Wnt pathway mutations that downregulate DC-attractant CCL3/4 chemokines (Spranger et al., 2015). However, most cancers do not carry these mutations. Additionally, Batf3-DC intratumoral paucity and dysfunction has been attributed to tumor-intrinsic disabling of Batf3-DC/ NK crosstalk through FLT3L and chemokines CCL5 and XCL1 (Barry et al., 2018; Bottcher et al., 2018). However, our data highlight a distinct mechanism through GM-CSF.

Activated Batf3-DC are preferentially located in the peritumoral matrix, distal to digitating tumor nestlets, where they are poised to interact with transiting CD8+ effectors (Broz et al., 2014; Hubert et al., 2020; Mattiuz et al., 2021). The accumulation and retention of immunogenic DC along the tumor rim presents a paradox that cannot easily be explained through simply invoking Batf3-DC displacement from tumor nests. The existing literature does not adequately explain why Batf3-DC are preferentially retained within peritumoral stroma and how they become “poised” to interact with T cells. Indeed, the mere maintenance of subsets of stimulatory Batf3-DC in tumors has been seen as “counterintuitive” (Balan et al., 2020). Therefore, we took a tumor-extrinsic viewpoint and reasoned that the accumulation, persistence and function of stimulatory DC at the tumor periphery may be dependent on invasive margin matrix remodeling signals. In embryonic development, non-redundant homeostatic signals arising from provisional matrix remodeling regulate cellular fates in the vicinity of tissue plane forging. This consideration brings into sharp focus matrix proteoglycans, key players in tissue sculpting in both embryonic and adult tissues. One of the most central actors in this group is versican (VCAN) (reviewed in (Papadas and Asimakopoulos, 2020)). VCAN proteolytic processing has been shown to be absolutely essential in crucial contexts including regression of interdigital webs, palate sculpting, myocardial compaction and heart valve morphogenesis (Nandadasa et al., 2014).

Our findings suggest that homeostatic signals emanating from stromal invasion support Batf3-DC survival and activity at the tumor edge. Stroma-licensed peritumoral Batf3-DC promote T-cell infiltration into tumor nests through re-priming and expansion of transiting effectors along invasive margins. In small nascent tumors, the process may well end in tumor elimination through Batf3- DC activation. Tumors that survive with active stromal-DC signaling may display T-cell inflammation. Attenuation of stromal DC-activating signals and/or enhancement of stromal DC- inhibitory cues redresses the balance and results in effector stalling within the tumoral border and “immune exclusion”. We illustrate these principles through the lens of the VCAN pathway: VCAN proteolysis to generate versikine represents a host homeostatic response to the expanding tumor. Versikine supports the survival and activation of peritumoral Batf3-DC and promotes a robust adaptive response and T-cell infiltration. Progressive immunosuppression results in attenuation of VCAN proteolysis, versikine depletion and accumulation of parental VCAN that disables peritumoral DC, resulting in immune exclusion, an effect that could be mediated through tolerogenic VCAN-TLR2 signaling (Tang et al., 2015) (Fig. 7G). Interestingly, TGF*β*, a known instigator of T-cell exclusion (Mariathasan et al., 2018), upregulates VCAN production and downregulates ADAMTS-cleaving versicanases (Cross et al., 2005).

There is concrete evidence supporting this model: non-proteolyzed VCAN has been linked to T-cell exclusion in human cancers ((Evanko et al., 2012; Gorter et al., 2010; McMahon et al., 2016) and Fig. 1 data in this manuscript). Conversely, in colorectal cancer, the highest intra-epithelial T-cell infiltration was associated with the VCAN-proteolysis-predominant phenotype, characterized by the combination of high stromal versikine and low non-proteolyzed VCAN (Hope et al., 2017).

The fact that versikine was insufficient to induce tumor regression on its own was somewhat surprising. However, our tumor models consisted of terminally immune-edited implantable cancers whose elimination may not be achieved through an increase in Batf3-DC density/ activation alone. Moreover, versikine promoted a “T-cell inflamed” phenotype that was still compatible with tumor propagation but at the cost of heightened sensitivity to physiological immunosurveillance mechanisms, e.g., nucleic acid sensing. The mechanistic basis for this DNA- hypersensitive phenotype could be related to Irf7 induction (Fig. 5). Indeed, IRF7 is a target of STING-mediated TBK1 induction through phosphorylation and subsequent activation of type I IFN transcription (Dalskov et al., 2020). Alternatively or additionally, versikine-mobilized T cells may be inhibited by immune checkpoints, e.g., PD-1, within the tumor mass.

The prevailing view in the literature casts peritumoral stroma overwhelmingly in the role of immune “barrier” (Joyce and Fearon, 2015) but our data support a more dynamic and fluid perspective. Transition towards a more nuanced distinction between “stimulatory stroma” and “regulatory stroma” may permit the design of more accurate personalized immunotherapy approaches. Early evidence from the clinic demonstrates superior responses to checkpoint inhibition by tumors displaying robust endogenous VCAN proteolysis (Deming et al., 2020). Importantly, harnessing stromal signaling can pave the way for therapeutic advances. Therapeutic repletion and redistribution of matrix-DC activation signals to the tumor core may be exploited to generate or potentiate a “hot” immune microenvironment that sensitizes immunoevasive tumors to immunotherapy, to promote antitumor responses and/or overcome resistance. Several approaches can be envisioned, including direct delivery into the tumor bed, or expression by engineered effectors or oncotropic vectors. Therapeutic recasting of tumor architecture could thus boost active or passive immunotherapy outcomes, particularly in the challenging settings of advanced or metastatic solid cancers.

## METHODS

### Animal strains and regulatory approval

C57BL/6J (JAX stock 000664), BALB/cJ (JAX stock 000651), B6.129 (Cg)-*Cd44^tm1Hbg^*/J (*Cd44*-/-, JAX stock 005085), B6.129S(C)-*Batf3^tm1Kmm^*/J (*Batf3*-/-, JAX stock 013755), B6.129-*Tlr2^tm1Kir^*/J (*Tlr2*-/-, JAX stock 004650), C57BL/6-Tg(TcraTcrb)1100Mjb/J (OT-I, JAX stock 003831) and VQ mice (Wen et al., 2021) were housed, cared for, and used in accordance with the *Guide for Care and Use of Laboratory Animals (NIH Publication 86-23)* under IACUC-approved protocols #M5476 and #S19109 in the University of Wisconsin-Madison and University of California, San Diego respectively.

### Generation of *Vcan* +/- mice using CRISPR-Cas9 gene editing

A mixture of two gRNA (25ng each, sgRNA#1 5’- ACTAGCCCGGAGTTTGACCA-3’, sgRNA#2 5’- ACCGATGTGATGTCATGTAT-3’) targeting mouse Vcan exon 3 and Cas9 protein (40ng; PNABio) was injected into the pronucleus of one-cell fertilized embryos from C57LB/6J females. Injected embryos were transferred into pseudo-pregnant females. Tail samples were taken at weaning, and the targeted region was characterized using targeted ultradeep sequencing. Briefly, the targeted region was PCR amplified using the following primers:

>207A.VCAN.ex2.F.1.6N.ILTS.1

acactctttccctacacgacgctcttccgatctNNNNNNACTGTCTTGGTGGCCCAGAAC

>207A.VCAN.ex2.R.1.ILTS.1

gtgactggagttcagacgtgtgctcttccgatctTCTCTGGTACCATGCTGCCTTTC

Samples were indexed & pooled, and the pool was sequenced on a MiSeq 2x250 Nano. Resultant sequences were quality filtered, trimmed, and analyzed with CRISPResso (Pinello et al, 2016). Founders were backcrossed to C57LB/6J mates, and F1s were characterized similarly.

For genotyping, DNA was extracted from mouse tail using genomic DNA extraction kit (Promega Wizard SV Genomic DNA Purification System, Catalog #: A2360), according to the manufacturer’s protocol. For PCR, Promega 2X GoTaq Master Mix (Catalog #: M7123), 1ul of template DNA and 10uM of each primer were used. The PCR conditions were 1min at 95°C followed by 35 cycles of 15 s at 95°C, 15 s at 60°C, 30 s at 72°C and a final extension of 1 min at 72°C. Primers for target sequences are listed on Supp. Table S7.

### Cell lines and primary cell culture

Lewis Lung Carcinoma (LLC, ATCC CRL-1642) and B16F10 melanoma (ATCC CRL-6322) were cultured in complete DMEM medium (10-013 CV Corning DMEM with 10% fetal calf serum, 50μM 2-mercaptoethanol, 100U/ml Penicillin, 100μg/ml Streptomycin, 292ng/ml L-Glutamine). 4T1 breast carcinoma cell line (CRL-2359) was cultured in complete RPMI (10-040 CV Corning RPMI 1640 with 10% fetal calf serum, 50μM 2-mercaptoethanol, 100U/ml Penicillin, 100μg/ml streptomycin, 10mM non-essential amino acids and 1M HEPES buffer). VQ4935 cells (Wen et al., 2021) were cultured in suspension in Iscove’s DMEM medium [10-016-CV Corning Iscove’s DMEM supplemented with 10% Fetal calf serum, 50μM 2-mercaptoethanol, 100U/ml penicillin, 100μg/ml streptomycin, 10mM non-essential amino acids and 10ng/mL IL-6 (Peprotech)]. Immortalized mouse dendritic cells (MutuDC1940, Applied Biological Materials Inc. #T0528) were cultured in Iscove’s DMEM medium supplemented with 10% fetal calf serum, 292ng/ml L-glutamine, 50μM 2-mercaptoethanol, 1% of 7.5% sodium bicarbonate (w/v). iCD103 *in vitro* differentiation was performed using bone marrow cells from female C57BL/6 mice at 6- 12 weeks of age according to the protocol by Mayer and colleagues (Mayer et al., 2014). 15- 16 days after start of the culture, DC were harvested, immunophenotyped and used for experiments.

### Constructs

pLenti6-UbC-VKine-HA and pLenti6-UbC-VKine-Myc has been previously described (Hope et al., 2016; McCulloch et al., 2009). Ovalbumin (OVA) amplicon was PCR amplified from pcDNA3-OVA (Addgene #64599) and cloned into pHIV-Luc-ZsGreen backbone (Addgene #39196). All lentiviral constructs were transformed into NEB 5-alpha competent cells (#C2987U) for propagation of plasmid DNA. All plasmids were prepped and purified using Macharey-Nagel NucleoBond Xtra Maxi kit (# 740414.50).

### Lentiviral transduction

HEK293T cells were transfected with a mixture of ps-PAX2 (packaging plasmid) and pVSV-G (envelope plasmid), and transfer plasmids encoding respective open reading frames or empty control. On Day 2 post-transfection, pseudotype virus-containing culture medium was harvested, filtered, supplemented with 7.5 μg/ml polybrene (Sigma-Aldrich), and immediately applied to target cells for spinfection (120min, 2500xg at 32C). After spinfection, the medium was exchanged for fresh complete RPMI1640 medium. Target cells were passaged at least three times after retroviral transduction.

### Generation of HA-tagged Versikine- and OVA-ZsGreen-expressing cell lines

LLC, 4T1, B16F10 melanoma and MutuDC1940 cells were transduced with HA tagged VKine- or empty vector (EV)- containing lentivirus as detailed above. The cells were selected with 10µg/ml blasticidin for 2 weeks. HA-tagged versikine expression was confirmed by western blotting using anti-HA antibody. LLC- EV or -Vkine cell lines were transduced with pHIV-Luc-OVA-ZsGreen lentivirus. LLC-OVA expressing cells were FACS-sorted based on ZsGreen expression to ensure comparable transduction rates between different cell lines.

### shRNA mediated VCAN knockdown

The lentiviral shRNA vector set targeting mouse Vcan (NM_019389.2) and scrambled control were purchased from GeneCopoeia (#MSH080253-LVRU6H and #CSHCTR001-LVRU6H). In brief, 2×10^5^ LLC cells were plated per well in a 6-well plate and incubated overnight. Next day, 2ml freshly harvested lentiviral supernatant (expressing either shambled control, Vcan shRNA#1, #2 or #3), 1 ml of culture medium and 7.5μg/ml polybrene was added per well. The plate was centrifuged at 800g for 2h at 37°C and returned to CO2 incubator. After 72h, 200μg/ml Hygromycin B was added and the cells were under antibiotic selection for 2 weeks. Vcan knockdown was confirmed by RT-PCR as shown in Supp. Fig. S1.

### Tumor cell inoculations and tumor growth measurement

Cells were harvested by trypsinization and washed in PBS. Mice were under isoflurane anesthesia during tumor injections. 5x10^5^ LLC cells were injected subcutaneously (s.c.) in 100μl endotoxin-free PBS on the flank of recipient mice. 10^5^ 4T1 cells were injected orthotopically in the mammary fat pad of the mice. Tumor growth was measured using a digital caliper. Tumor volumes were measured bi-weekly and estimated by using the formula: Tumor volume= length x (width) ^2^ divided by 2, where length represented the largest tumor diameter and width represented the perpendicular tumor diameter. Intratumoral injections were performed using a 28G insuilin syringe, when tumors had reached 100-150 mm^3^, using surgical forceps to hold the tumor constantly. For intravenous (i.v) incoculations, we adopted a retro-orbital approach. Mice were anesthetized using inhaled isoflurane in a chamber. The eyeball was partially protruded from the socket by applying downward pressure to the skin dorsal and ventral to the eye. Injections were performed by placing the needle, so the bevel faced down, in order to decrease the likelihood of damaging the eyeball. Once the injection was complete, the needle was slowly and smoothly withdrawn. Triple antibiotic ophthalmic ointment was then applied to the eye.

Intraperitoneal injection was performed using a 28.5G insulin-syringe with the head tilted down. The needle was inserted at a 30° angle in the lower left or right quadrant. Transplantation of myeloma VQ4935 cells was performed via intracardiac injection after the 6-8 week old C57BL/6J recipient mice were sub-lethally irradiated at 6.0 Gy using an X-RAD 320 Irradiator. Intracardiac injection was performed by placement of needle in the 4^th^ intercostal space and into the left ventricle. The needle was inserted at a 90° angle in the middle of the imaginary line connecting the sternal notch and xyphoid process serving as anatomical landmarks, and the needle was inserted slightly left of the sternum.

### Processing of tumor tissue

Unless stated otherwise, tumors were excised 21 days after transplantation. For subsequent analysis by flow cytometry, tumors were cut into pieces and digested with Collagenase Ia (1mg/ml) C2674 Sigma Aldrich and Hyaluronidase V (0.1mg/ml) H6254 Sigma Aldrich for 40min at 37**°**C using gentle MACS dissociator. Tissue was passed through a 70μm cell strainer (Falcon) and washed with FACS buffer (PBS with 1% FCS) before proceeding with antibody staining. For RNA isolation, homogenization was performed in RLT buffer (QIAGEN) facilitated by a closed tissue grinder system (Fischer brand #02-542-09, 15mL).

### Mass cytometry

Tumor tissue was harvested and processed for mass cytometry analyses using the protocol described above for flow cytometry. After single cell suspensions were acquired, cells were washed with PBS, centrifuged at 300-400g for 5 minutes and supernatant was discarded by aspiration. Cells were resuspended in PBS and Cell-ID Cisplatin (Fluidigm, #201064) was added to a concentration of 5uM. After rigorous mixing, cells were incubated at room temperature for 5 minutes. Cells were then quench stained with MaxPar Cell Staining Buffer (Fluidigm, #201068) using 5x the volume of the cell suspension, centrifuged and supernatant was discarded by aspiration. The process was continued with surface staining. 50ul of the antibody cocktail was added to each tube so the total staining volume was 100ul (50ul of cell suspension+ 50ul antibody cocktail). Cells were stained for an hour at room temperature. All antibodies used for staining were either bought pre-conjugated to metal isotopes or were conjugated using the Maxpar Antibody Labelling Kit (Fluidigm 201160B) (Supplementary Table S7). Following incubation, cells were washed by adding 2mL Maxpar Cell Staining Buffer to each tube, then centrifuged at 300xg for 5 minutes and supernatant was removed by aspiration. This step was repeated for a total of 2 washes, and cells were resuspended in residual volume by gently vortexing after final wash/aspiration. Cells were then fixed with 1.6% FA solution and incubated at room temperature for 10 minutes. Finally, cells were labelled with Cell-ID Intercalator-Ir (Fluidigm, #201192A) at a final concentration of 125nM, incubated for an hour at room temperature and then analyzed on a Helios instrument (WB injector). All samples were resuspended in sufficient volume of 0.1 EQ beads (Fluidigm, #201078 by diluting one part beads to 9 parts Maxpar Cell Acquisition (CAS) solution.

### Analysis of mass cytometry data using viSNE

To visualize the immune contexture, the immune milieu of the tumor (CD45+) was enriched by manual gating among single events, equally subsampled to 6,000 events, then run through a Barnes Hut implementation of the t-SNE algorithm, viSNE, in the R package ‘Rtsne’, using optimized parameters (iterations:1000, perplexity:30, learning rate:455). All markers listed in Table S2 to characterize the myeloid and lymphoid linages were selected for viSNE, excluding CD45.

### Flow cytometry and fluorescence activated cell sorting

Flow cytometric analyses were performed using an LSR II and/or LSR Fortessa X20. Data were analyzed using FlowJo (Tree Star). DAPI (0.5 mg/ml, Sigma-Aldrich) or a Live/Dead fixable cell stain (Ghost 780 Tonbo Biosciences) was used to exclude dead cells in all experiments, and anti- CD16/CD32 antibody (2.4G2) was used to block non-specific binding of antibodies via Fc-receptors. The following antibodies were used for flow cytometry: anti-CD24 (clone M1/69), anti- CD45 (30-F11), anti-CD11c (N418), anti-CD11b (M1/70), anti-CD64(X54-5/7.1), anti-Ly6C (HK1.4), anti-CD103(2E7), anti-MHC II I-a/I-E (M5/114.15.2), anti-CD135 (A2F10), anti-CD172a (SIRPa) (P84), anti-CD45.1 (A220) anti-CD45.2 (104), anti-Clec9a/DNGR-1 (7H11), anti-NK1.1 (PK136), anti-CD49b(DX5), anti-CD3e (145/2C11), anti-NKp46 (29A1.4), anti-CD3 (17A2), anti- IFN*γ* (XMG1.2), anti-CD8*α* (53-6.7), anti-IL-2 (JES6-5H4), anti- H-2k(b) SINFEKL, anti-CD44 (IM7), anti-CD62L (MEL-14), anti-CD138 (syndecan-1) (281-2). NK cells were identified as live CD45^+^NK1.1^+^CD49b^+^CD3^−^MHCII^−^ cells. CD103^+^ cDC1 were identified as live CD45^+^ Cd24^+^CD103^+^CD11b^−^CD11c^+^ MHCII^+^ cells. Quantification of total cell numbers by flow cytometry was done using fluorescent beads (Biolegend Precision beads). For intracellular staining of IFN*γ* and IL-2 *in vitro*, cells were treated with Golgi Plug (Brefeldin A 500x) and were collected 4h later. Intracellular staining was performed in permeabilization buffer (eBioscience) for 30min and cells were subsequently analyzed by flow cytometry. Antibodies were purchased from Biolegend or BD Biosciences, as shown. Sorting of tumor cells after retroviral transduction was done using a BD FACSAria or a BD FACSAria Fusion. Purity of cell populations was determined by re-analysis of a fraction of sorted cell samples.

### Generation of iCD103 *in vitro*

For the generation of iCD103, 1.5x10^6^ BM cells were cultured in 10ml RPMI1640 medium supplemented with 10% heat-inactivated FCS (Biochrom), penicillin/streptomycin and 50μM β- mercaptoethanol. Recombinant human Flt3L (300-19, Peprotech) and recombinant murine GM-CSF (315-03, Peprotech) were added at D0 of the culture. 5ml complete medium was added between D5 and D6 to minimize apoptosis. Non-adherent cells were harvested on D9, counted and re-plated at 3x10^6^cells in 10ml complete medium supplemented with Flt3L and GMCSF as on D0. Non-adherent iCD103 were harvested on D15-16. Cells were then validated by assaying for CD103, CD24, Clec9A, and CD11c by flow cytometry.

### ELISA

MutuDC1940 cells were left unstimulated or were in vitro stimulated with LPS for 8 or 24 hours at 37°. Cell-free supernatant was assessed for CXCL9 (R&D Quantikine mouse CXCL9 #MCX900) and IL27p28 (R&D Quantikine mouse IL27p28 #M2728) protein levels by ELISA according to the manufacturer’s instructions (R&D). For the antigen-presentation assay, cell-free supernatants were collected and assessed for IFN*γ* levels (R&D Quantikine mouse IFN*γ* #P233156).

### Immunoblotting

Whole-cell lysates were prepared by boiling cells in Laemmli Sample Buffer (Bio-Rad) supplemented with 100 mM DTT for 10 min at a final concentration of 10^7^ cells per milliliter. A total of 10^5^ cells or 20 mg protein was resolved by SDS-PAGE and transferred to Immobilon-P PVDF membranes (Millipore). Membranes were blocked in 5% milk in TBS-T (25 mM Tris-HCl [pH 7.4], 0.13 M NaCl, 2.7 mM KCl). Primary antibodies (anti-HA [C29F4; Cell Signaling Technologies], anti-DPEAAE [PA1-1748A; Thermo]) were diluted in 5% milk–TBS-T, and membranes were incubated overnight at 4 °C. Secondary Ab–HRP conjugate, as well as anti- GAPDH– HRP conjugate (A00192; GenScript), incubations were carried out for 1 h at room temperature. Signal detection was achieved using Amersham ECL.

### Immunohistochemistry

Paraffin-embedded murine tumor sections and unstained 4-5 μm-thick human lung carcinoma TMA (US Biomax Inc., BC041115e) sections were deparaffinized and rehydrated using standard methods. Antigen retrieval was carried out in citrate buffer, pH 6.0 (Vector Laboratories, #H-3300) for DPEAAE and HA; and pH 8.0 for XCR1 and CD8 (Abcam, ab93680). Primary antibodies included αDPEAAE (PA1-1748A, Thermo Fisher), anti-HA (C29F4, Cell Signaling Technology), anti-XCR1 (D2F8T, Cell Signaling Technology) and anti-CD8 (C8/144B, Ebioscience). The αDPEAAE neoepitope antibody has been previously validated (Foulcer et al, 2015). Stained slides were examined using an Echo Revolve microscope with attached digital camera. αDPEAAE immunostaining score was assessed by scoring staining intensity (0 for no staining, 1 for low/weak staining, 2 for moderate staining and 3 for strong/intense staining) as previously described (Hope et al., 2017).

### RNA isolation and quantitative real-time PCR

RNA was isolated using QIAGEN RNeasy Mini Kit and cDNA was synthesized using the iScript Reverse Transcription Supermix (Biorad). Quantitative real-time (qRT-PCR) analysis was performed using SsoAdvanced Universal SYBR Green Supermix (Biorad) according to the manufacturer’s instructions on an CFX96 Touch Real Time PCR detection (Biorad) using the relative standard curve method. PCR conditions were 2min at 50°C, 10min at 95°C followed by 40 2-step cycles of 15 s at 95°C and 1 min at 60°C. Primers for the targets listed in Supp. Table S7 as well as SDHA for normalization control, were used to assess relative gene expression.

For DMXAA-response analysis, RT² Profiler PCR Array (QIAGEN, Cat. no. PAMM-021Z) was used. In brief, RNA was isolated from tumors and was reverse transcribed using kits mentioned above. cDNA was mixed with RT² SYBR® Green qPCR Mastermix (Cat. no. 330529). The mixture was aliquoted across the RT² Profiler PCR Array (in 96-well format) and was run on the Real Time PCR machine. Data analysis was performed using the manufacturer’s online platform for RT² Profiler Data analysis software.

### Generation of versikine cell culture supernatant

5x10^6^ HEK293 and HEK-Vkine expressing cells (a kind gift of Dr. Suneel S. Apte, Cleveland Clinic Lerner Research Institute) were seeded in T-175 cell culture flasks and cultured in DMEM 10% FBS media. After 75 to 80% confluency, the cell media was changed to DMEM 1% FBS media. Subsequently, media supernatant was collected after 48 hours of incubation. The collected supernatant was centrifuged to remove debris and filtered with 0.45µ filter. The filtered supernatant was then concentrated 30 times to the initial volume using Sartorius Vivaspin 20, 10,000 MWCO PES concentrator (Cat. No. VS2001). Endotoxin assay was performed using Genescript ToxinSensor Gel Clot Endotoxin Assay Kit (Cat. No. L00351) according to the manufacturer’s instructions to rule out contamination. The presence of versikine in concentrated supernatant was confirmed using western blot using c-Myc Antibody (Novus Bio- c-Myc Antibody (9E10) - Chimeric NBP2-52636). Concentrated supernatant containing versikine was then used to treat MutuDC1940 cells. 2 x10^5^ MutuDC1940 cells per well were seeded in 12 well plate. The following day, 10% supernatant was added to the plate media. Cells were then incubated for 72 hours. After incubation, total RNA was extracted from MutuDC1940 cells.

### Library preparation for RNA-seq

A total amount of 1 μg RNA per sample was used as input material for the RNA sample preparations. Sequencing libraries were generated using NEBNext® UltraTM RNA Library Prep Kit for Illumina® (NEB, USA) following manufacturer’s recommendations and index codes were added to attribute sequences to each sample. Briefly, mRNA was purified from total RNA using poly-T oligo-attached magnetic beads. Fragmentation was carried out using divalent cations under elevated temperature in NEBNext First Strand Synthesis Reaction Buffer (5X). First strand cDNA was synthesized using random hexamer primer and M-MuLV Reverse Transcriptase (RNase H-). Second strand cDNA synthesis was subsequently performed using DNA polymerase I and RNase H. Remaining overhangs were converted into blunt ends via exonuclease/polymerase activities. After adenylation of 3’ ends of DNA fragments, NEBNext Adaptor with hairpin loop structure were ligated to prepare for hybridization. In order to select cDNA fragments of preferentially 150∼200 bp in length, the library fragments were purified with AMPure XP system (Beckman Coulter, Beverly, USA). Then 3 μl USER Enzyme (NEB, USA) was used with size-selected, adaptor-ligated cDNA at 37 °C for 15 min followed by 5 min at 95 °C before PCR. Then PCR was performed with Phusion High-Fidelity DNA polymerase, Universal PCR primers and Index (X) Primer. At last, PCR products were purified (AMPure XP system) and library quality was assessed on the Agilent Bioanalyzer 2100 system. The clustering of the index-coded samples was performed on a cBot Cluster Generation System using PE Cluster Kit cBot-HS (Illumina) according to the manufacturer’s instructions. After cluster generation, the library preparations were sequenced on an Illumina platform and paired-end reads were generated.

### RNA-seq data analysis

Raw data (raw reads) of FASTQ format were firstly processed through fastp. In this step, clean data (clean reads) were obtained by removing reads containing adapter and poly-N sequences and reads with low quality from raw data. At the same time, Q20, Q30 and GC content of the clean data were calculated. All the downstream analyses were based on the clean data with high quality. Reference genome and gene model annotation files (GRCm38) were downloaded from genome website browser (NCBI/UCSC/Ensembl) directly. Paired-end clean reads were aligned to the reference genome using the Spliced Transcripts Alignment to a Reference (STAR) software (v2.6.1d). FeatureCounts (v1.5.0-p3) was used to count the read numbers mapped of each gene. RPKM of each gene was calculated based on the length of the gene and reads count mapped to this gene. Differential expression analysis between two conditions/groups (three biological replicates per condition) was performed using DESeq2 R package (v1.20.0). DESeq2 provides statistical routines for determining differential expression in digital gene expression data using a model based on the negative binomial distribution. The resulting P values were adjusted using the Benjamini and Hochberg’s approach for controlling the False Discovery Rate (FDR). Genes with an adjusted P value < 0.05 found by DESeq2 were assigned as differentially expressed.

### Gene Set Enrichment Analysis

Gene Set Enrichment Analysis (Subramanian et al., 2005) was performed by comparing MutuDC1940-VKine (treated with PBS, 4h) RNA-seq data to the corresponding MutuDC1940-EV sample. 4736 differentially expressed gene features for each condition were ranked by the signal to noise metric of GSEA and the analysis was performed using the standard weighted enrichment statistic against human gene sets contained in the Molecular Signatures Database (MSigDB v7.4) that included all (H) Hallmark gene sets, (C2) curated gene sets, and (C3) motif gene sets. The normalized enrichment score (NES) was calculated using 1000 gene set permutations.

### NK cell depletion *in vivo*

For depletion of NK cells, mice were injected i.p. with 50 ug of anti-asialoGM1 (Wako Pure Chemical Industries, 100μl/mouse) on days -1, 0, 7, 14 post tumor inoculation.

### Antigen presentation assay

MutuDC1940 were cultured and treated with LPS or PBS control respectively overnight. Next day, the cells were harvested and plated on 96-well round bottom plates at a density of 100,000 cells per plate. DC were then loaded with OVA peptide 257-264 (SIINFEKL) (3ng/ml) and incubated for 4 hours at 37°C. MutuDC were then washed with 0.1% PBS-BSA and centrifuged at 800 x g and were fixed with 50μl per well of freshly made PBS-glutaraldehyde (GTA) 0.008% (vol/vol) and incubated for 5 minutes on ice. 50μl of PBS-glycine 0.4M was added to the PBS-GTA 0.008% solution and cells were centrifuged at 800 x g for 2 minutes at 4°C. Plates were subsequently flicked. Finally, 100 μl of PBS-glycine 0.2M was added to each well and centrifugation of the plates at 800 x g for 2 min at 4°C followed. Fixed DC were then washed twice with 200 μl/well of T-cell culture medium (RPMI 1640 containing 10% heat inactivated FBS, 100 IU/ml penicillin, 100μg/ml streptomycin, 2mM glutamax, 50μM *β*-mercaptoethanol, 1xMEM non-essential amino acids, 1x sodium pyruvate) before being resuspended in 100 ul/well of the same medium. 100,000 OT-I T cells per well in 100μl T cell culture medium were added (to a final volume of 200μl). The co-cultured OT-I T cells with the cross-fixed DCs were incubated for 18 h at 37°C. Cell activation cocktail with brefeldin A (PMA/Ionomycin and Brefeldin A Biolegend, #423303) was added to the wells 4 hours before harvesting. At the time of the harvest, plates were spun down at 800 x g for 2 minutes at 4°C and supernatant was kept for subsequent cytokine analysis. They were then washed with 100μl of 0.1% PBS-BSA before proceeding with live/dead staining with fixable viability Ghost 780 dye (Tonbo Biosciences) for 30 min at 4°C in PBS. Cells were then washed with 0.1% PBS-BSA and stained with a cocktail of 70μl/well of the surface markers (CD8*α* and CD3) for 40 minutes at 4°C. They were then fixed and permeabilized using the eBiosciences fixation and permeabilization buffer set (eBioscience 88-8824-00) according to the manufacturer’s instructions followed by intracellular staining of IFN*γ* and IL-2 in permeabilization buffer. Finally, cells were washed 2 times with 0.1% PBS-BSA, centrifuged and resuspended in 100μl/well of PBS-BSA and then analyzed by flow cytometry.

### Batf3-DC and VCAN landscape analysis of TCGA datasets

Level 4 gene expression data were downloaded from the TCGA Data Portal and filtered to retain only cancer types of known epithelial origin for a total of 7591 samples across 20 different cancer types. Single-sample Gene Set Enrichment Analysis (ssGSEA) was performed as described in Barbie and co-authors (Barbie et al., 2009) to measure the signature of gene sets designed to measure overall immune infiltration (Yoshihara et al., 2013), Batf3-DC density and CD8+ T cell density, as previously published (Spranger et al., 2017). TCGA samples were grouped by cancer type and sorted based on median expression of versican (VCAN) and median Batf3-DC signature. To measure cancer-specific relationships between VCAN expression and the Batf3-DC signature an ordinary least squares linear model was fit on these two variables to measure their relationship within each cancer type. Nominal p-values from these 20 different models were corrected for multiple hypothesis testing using the Benjamini-Hochberg method and q-values less than 0.1 were considered statistically significant.

### Computational modeling of CD8+ T cell density and versikine response signature

Differential expression using DESeq2 was performed to identify genes that were differentially expressed between the PBS-EV and PBS-versikine conditions (Fig. 4A and 4C). Genes with a q-value of less than 0.1 were considered significant. We define two gene sets to measure the response to versikine by selecting the 100 genes more significantly induced (versikine-up) and most significantly repressed (versikine-down). These mouse genes were then mapped to their human orthologues using the HGNC Comparison of Orthology Predictions (HCOP) tool (Eyre et al., 2007). ssGSEA (Barbie et al., 2009) was then used to measure the signature of these two gene sets in the 1017 lung samples in the TCGA cohort and overall versikine response level was summarized as the difference between versikine-up and versikine-down signature levels. This versikine response signature was then compared to the CD8 effector T cell signature using an ordinary least squares linear model including the overall immune infiltration signature as a covariate. P-values of less than 0.05 were considered significant.

### Graphics

Graphics and diagrams were created using BioRender and Omnigraffle.

### Statistical analysis

Statistical analysis was performed using GraphPad Prism software (GraphPad version 9.0.0) or Python version 3.6.6 with stats models version 0.10.0. Statistical significance was determined using unpaired two-tailed Student’s t-test or unpaired non-parametric Mann-Whitney test as indicated in figure legends. The log-rank (Mantel-Cox) test was used to determine statistical significance for overall survival in *in vivo* experiments. Data are shown as mean ± SD. Significance was assumed with *p<0.05; **p<0.01; ***p<0.001, ****p<0.0001.

## ACKNOWLEDGMENTS

This work was supported through the NIH/ National Cancer Institute (R01CA252937, R01CA230275 and U01CA196406), the American Cancer Society (127508-RSG-15-045-01-LIB), the Leukemia and Lymphoma Society (6551-18), the UW Trillium Myeloma Fund and the Robert J. Shillman Foundation. APapadas was supported through scholarships from the A.G. Leventis Foundation, the Gerondelis Foundation and the Mentzelopoulos Foundation. AO was supported by the US NLM (T15LM011271). APapadas and AO were recipients of AACR Scholar-In-Training awards and APapadas was recipient of a SITC Young Investigator award.

We thank Cheryl Kim, Denise Hinz, Chris Dillingham and staff at the La Jolla Institute of Immunology Flow Cytometry facility; Dagna Sheerar and staff at the UW Carbone Cancer Center Flow Cytometry facility; Jesus Olvera and Cody Fine at the UCSD Human Embryonic Stem Cell Core Facility; Dustin Rubinstein and Kathy Krentz at the UW Biotechnology Center; for help with flow cytometry, mass cytometry and transgenic animal generation, respectively. We thank Elsa Molina at the UCSD Stem Cell Genomics Facility for help with gene expression assays. We thank Suneel S. Apte (Lerner Research Institute at Cleveland Clinic) for the kind gift of versikine- expressing HEK293 cells. We thank Emanuela Romano, Marine Gros and Amigorena lab members (Institut Curie, Paris, France) for advice on antigen-presentation assays. We thank Damya Laoui (Free University, Brussels, Belgium) for advice on DC gating strategies.

We thank Gutkind lab members (UCSD) for moral support and sharing equipment. We thank Joe Aguilera at UCSD Moores Cancer Center for help with lab relocation management. We thank Sasha Rakhmilevich and Paul Sondel (both at UW-Madison) for advice and support. We thank the UW CMP Graduate Program, Zsuzsa Fabry, Joanne Thornton, Christian Capitini, Mario Otto and Peiman Hematti for career advice and support to APapadas’s graduate studies.

## AUTHOR CONTRIBUTIONS

APapadas, GD, AO, AC, CH, DD, KP, SIA, OH and FA designed experiments and collaborated to develop the rational structure of the study. APapadas, GD, AC, CH, PE, JW, APagenkopf, GA and VB performed experiments. KAM provided pathology expertise. AO and OH performed computational analysis. FA was overall responsible for design and conduct of the study and wrote the first draft of the manuscript. All co-authors reviewed, revised and approved the final manuscript.

## SUPPLEMENTARY FIGURE LEGENDS

**Supplementary Fig. S1 (related to Fig. 1).**
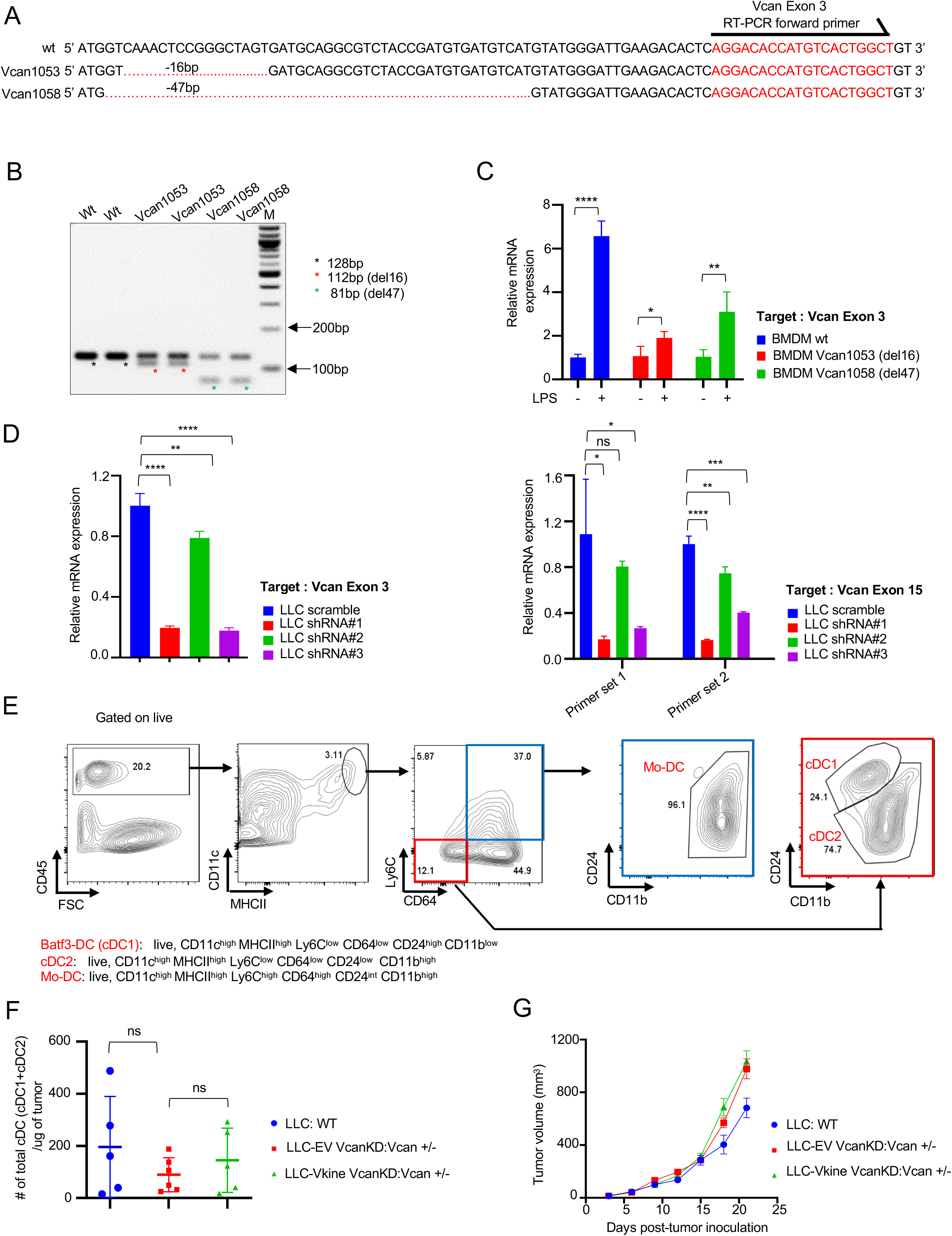
A: Schematic depiction of the deletions in the two mutant *Vcan* founders, 1053 (16bp deletion) and 1058 (47bp deletion). Sequence of exon 3 primer used in RT-PCR experiments shown in red. B: DNA amplification using primers flanking the targeted region. Shown are a 128bp WT amplicon and the mutated amplicons in founders *Vcan*1053 and *Vcan*1058. C: Bone marrow-derived macrophage (BMDM) *Vcan* locus RT-PCR using exon 3 primers at baseline or after stimulation with LPS. *Vcan*1053 demonstrates a more severe defect in *Vcan* message induction/stability than *Vcan*1058. D: Validation of *Vcan* knockdown in LLC^VcanKD^ cells. LLC cells were transfected with each of 3 hairpins (shRNA #1, 2 or 3) targeting exon 8 (encoding GAG*β* domain depicted in red in Fig. 1A). *Vcan* message was assayed using exon 3 primers (left) and exon 15 primers (right). E: Gating strategy to delineate tumor-associated dendritic cells (TADC) per Van Ginderachter schema (Laoui et al., 2016). F: Total cDC (cDC1 + cDC2) absolute counts per mcg tumor mass in WT, Vcan-depleted (Vcan+/-: LLC^VcanKD^-EV) and versikine-rescue (Vcan+/-: LLC^VcanKD^-Vkine) tumors. G: Growth rates of WT, Vcan-depleted (Vcan+/-: LLC^VcanKD^-EV) and versikine-rescue (Vcan+/-: LLC^VcanKD^-Vkine) tumors. Data are represented as mean±SD. n=5 or 6 for each group. *p<0.05; **p<0.01;***p<0.001.

**Supplementary Fig. S2 (related to Fig. 2).**
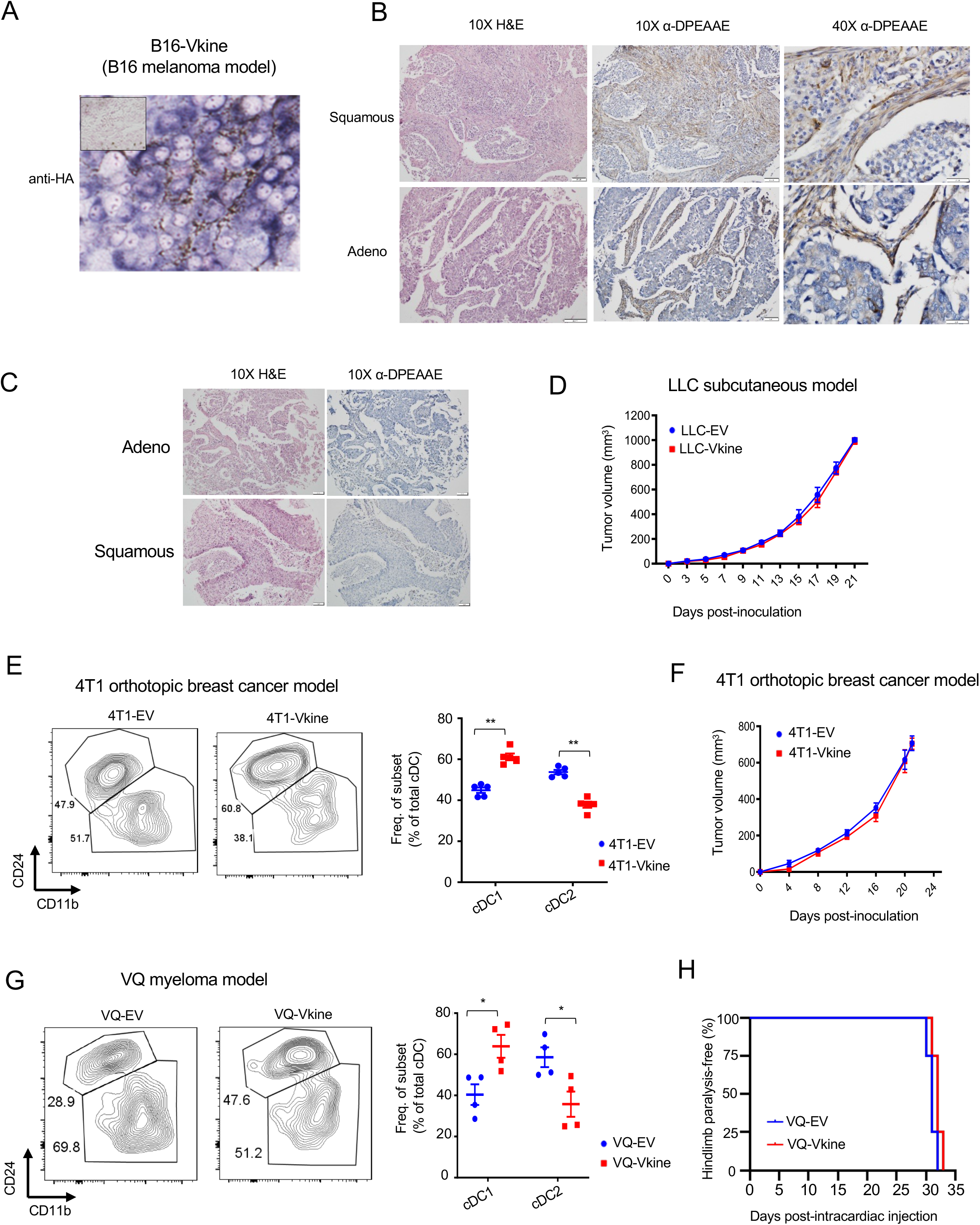
A: Membranous localization of HA-tagged ectopic versikine in a tumor model characterized by absence of cell-autonomous *Vcan* expression (B16 melanoma). HA stain, chromogen: BCIP/NBT; counterstain: nuclear fast red. B: Examples of human lung cancers with subpopulations of tumor cells staining positive for DPEAAE in an epithelial distribution. The vast majority of these cases also show concurrent stromal staining (Supp. Table S1). 10X objective: scalebar 50 μm, 40X objective: scalebar 20μm. C: Representative immunohistochemistry images showing examples of negative DPEAAE staining in major human lung cancer histologies. 10X objective: scalebar 50 μm. See Supp. Table S1 for summary of staining patterns. D: Growth rates of subcutaneous LLC-EV and LLC-Vkine tumors. E: Flow cytometric analysis of cDC subsets in 4T1 breast cancer cells orthotopically injected into the cleared fat pad of Balb/c recipient mice, engineered to express empty-vector (4T1-EV) or versikine (4T1-Vkine). Representative flow plots (top) and quantification (bottom) of cDC subsets are shown. F: Growth rates of orthotopic 4T1-EV and 4T1-Vkine tumors. G: Flow cytometric analysis of cDC subsets of tumors developed from intracardiac injection of VQ myeloma cells engineered to express empty-vector (VQ-EV) or versikine (VQ-Vkine). H: Kaplan-Meier curves depicting time-to-hindlimb paralysis (a clinical sequela of myeloma progression) in VQ-EV vs. VQ- Vkine myeloma tumors. In (E) and (G): *p<0.05; **p<0.01;***p<0.001.

**Supplementary Fig. S3 (related to Fig. 3).**
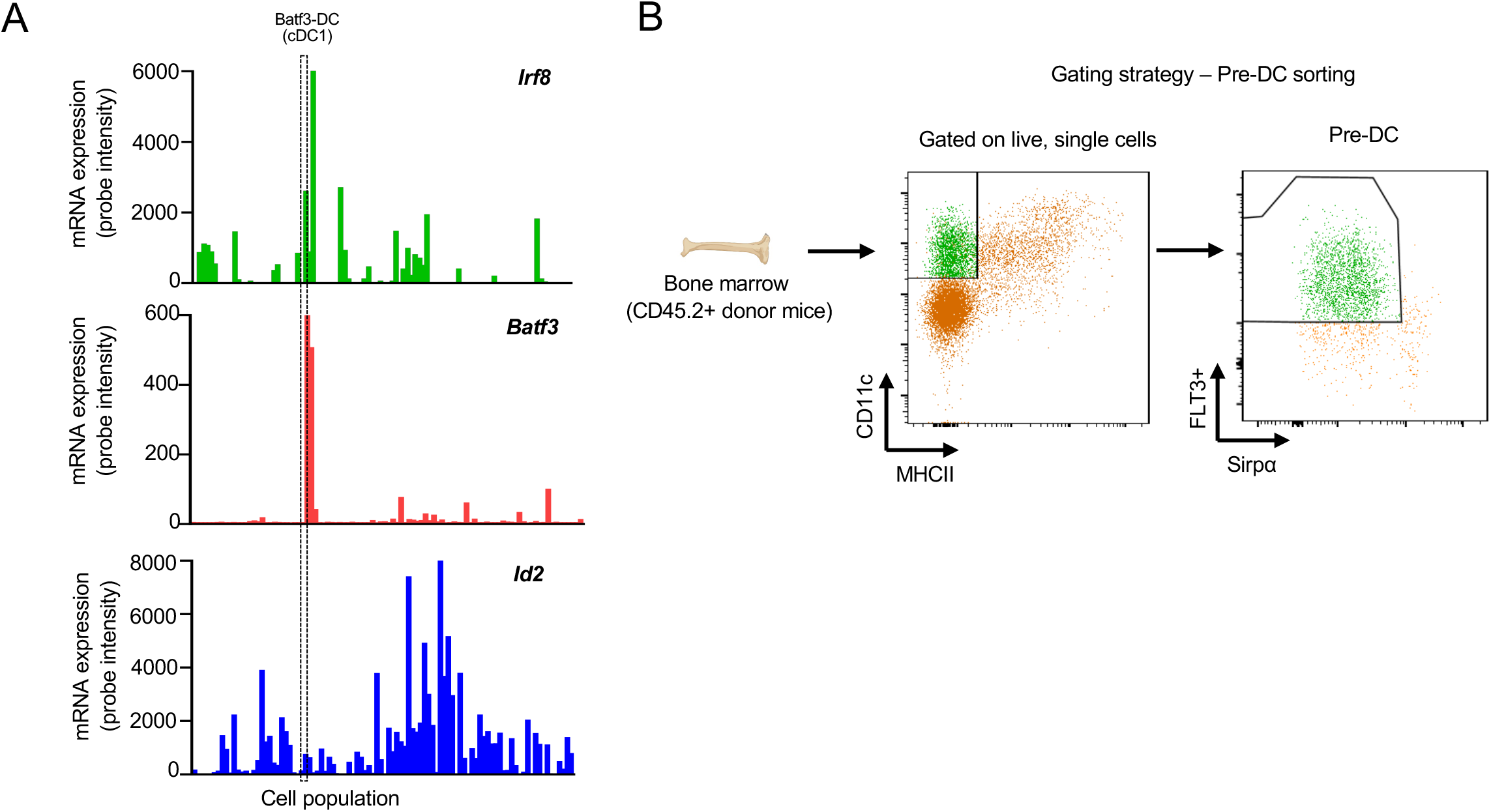
A: Expression pattern of murine *Irf8, Batf3 and Id2*. Data from BioGPS. See Table S3 for tissue/ lineage annotations. B: Gating strategy for flow sorting pre-DC from Flt3L-mobilized donors, per the schema of Van Ginderachter (Laoui et al., 2016). DC were mobilized *in vivo* in donor mice bearing Flt3L-secreting B16 tumor cells.

**Supplementary Fig. S4 (related to Fig. 4).**
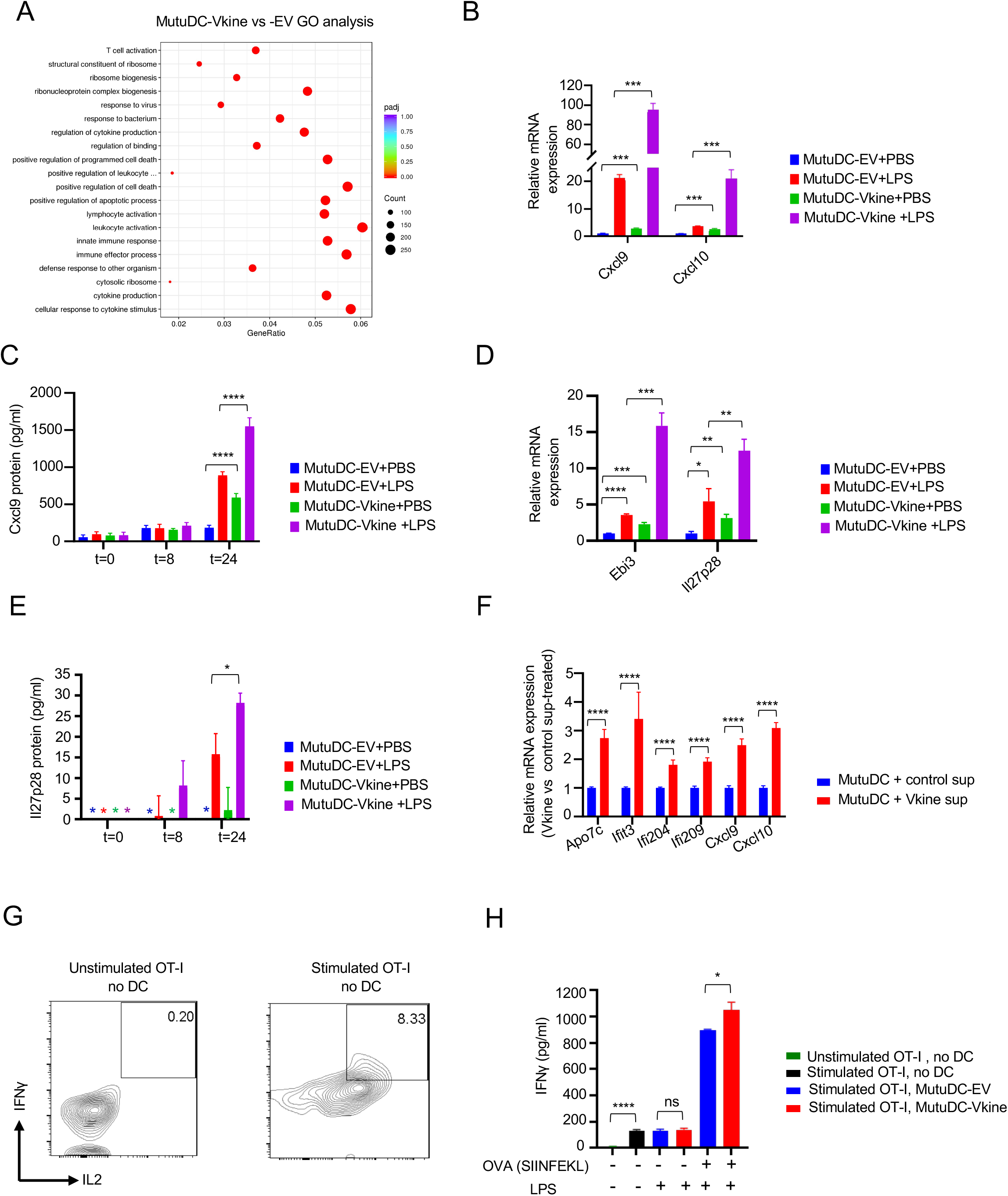
A: Gene ontology pathway analysis of differentially expressed genes between MutuDC1940-Vkine vs. -EV. Proapoptotic pathway annotation is reminiscent of versikine’s reported apoptotic, limb-sculpting activities during development (McCulloch et al., 2009). B: RT-PCR of Cxcl9/10 in MutuDC1940-EV vs. MutuDC1940-Vkine stimulated with LPS or vehicle (PBS). C: ELISA detection of secreted Cxcl9 by MutuDC1940-EV- and MutuDC1940-Vkine stimulated with LPS or vehicle (PBS) plotted against time (hours). D: RT-PCR for Il27p28 and Ebi3 message in MutuDC1940-EV-vs. -Vkine stimulated with LPS or vehicle (PBS). E: ELISA detection of secreted Il27p28 by MutuDC1940-EV- and MutuDC1940-Vkine stimulated with LPS or vehicle (PBS) plotted against time (hours). F: RT-PCR for selected versikine-signature genes using RNA from MutuDC1940 cells exposed to supernatant from versikine-secreting HEK293 cells (Vkine sup) vs. control supernatant (Control sup) at 72 hours. G: Flow cytometry for endogenous IFN*γ* and IL2 of OT-I CD8+ T cells at baseline (left) and PMA-stimulated, prior to addition of DC (right). H: IFN*γ* by ELISA in supernatants from OT-I+ MutuDC1940:SIINFEKL co-cultures in the antigen presentation assay. Data are represented as mean±SD, n=3, ns, non-significant, *p<0.05; **p<0.01;***p<0.001; ****p<0.0001.

**Supplementary Fig. S5 (related to Fig. 5).**
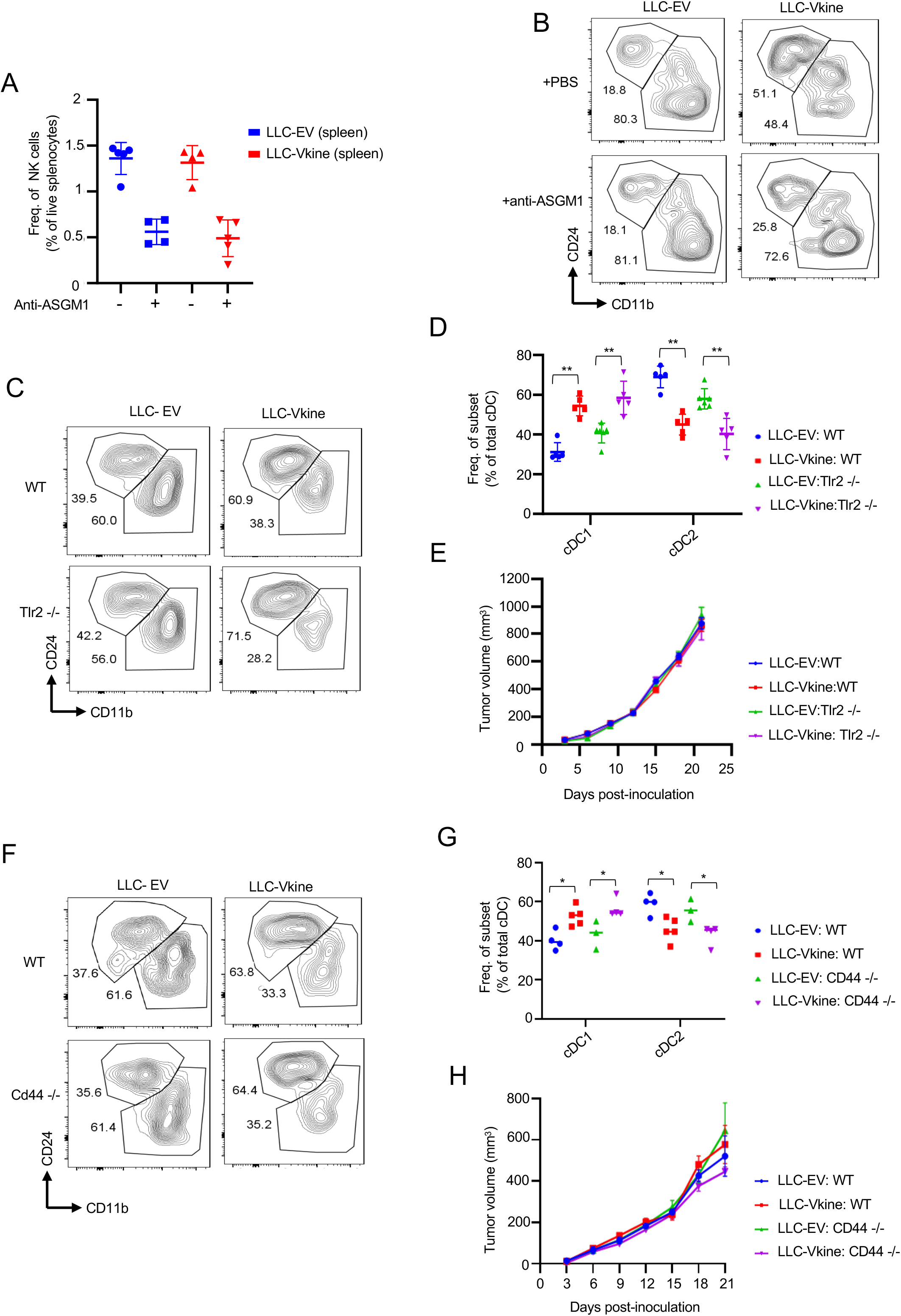
A: Validation of splenic NK depletion following anti- ASGM1 treatment. B: Flow cytometric analysis of cDC subsets in LLC-EV vs. LLC-Vkine tumors following treatment with NK-depleting antibody (anti-ASGM1) or vehicle (PBS). C: Flow cytometric analysis of cDC subsets in LLC-EV vs. LLC-Vkine tumors implanted in WT or *Tlr2-/-* recipients. D: Summary of cDC subset frequency by flow cytometric analysis in LLC-EV vs. LLC-Vkine tumors implanted in WT or *Tlr2-/-* recipients. E: Growth rates of LLC-EV and LLC-Vkine tumors in WT vs. *Tlr2*-/- background. F: Flow cytometric analysis of cDC subsets in LLC-EV vs. LLC-Vkine tumors implanted in WT or *Cd44-/-* recipients. G: Summary of cDC subset frequency by flow cytometric analysis in LLC-EV vs. LLC-Vkine tumors implanted in WT or *Cd44-/-* recipients. H: Growth rates of LLC-EV and LLC-Vkine tumors in WT vs. *Cd44*-/- genetic background.

**Supplementary Fig. S6 (related to Fig. 6).**
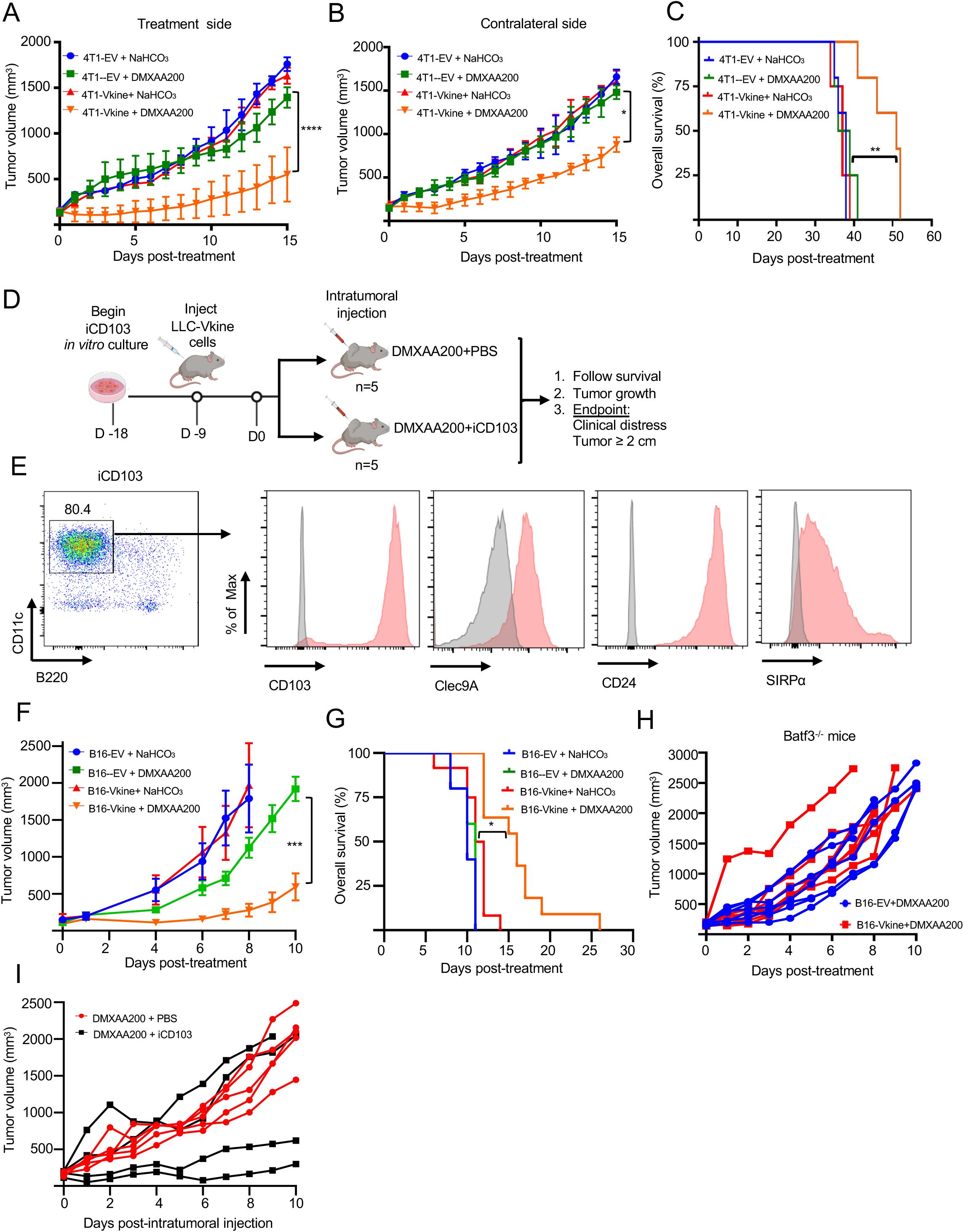
A: Versikine-DMXAA synergy generates an abscopal effect in 4T1 tumors. Growth curves of <u>treatment-side</u> 4T1-EV and 4T1-Vkine tumors challenged with a single subtherapeutic dose (200 mcg) of IT DMXAA on Day 0 (DMXAA200) or vehicle (NaHCO3). B: Growth curves of <u>contralateral side</u> unmanipulated 4T1 tumors, according to corresponding treatment side configuration (treatment as in Panel S6A). C: Versikine-induced abscopal effect is accompanied by a survival advantage in 4T1 tumors. **=p<0.01 by log-rank test. D: Schematic layout of iCD103 adoptive transfer experiments. E: Flow-cytometric validation of the iCD103 cells, generated as described in the protocol by Merad, Sparwasser and colleagues (Mayer et al., 2014), using standard Batf3-DC markers. F: Growth curves of B16-EV and B16- Vkine tumors challenged with a single subtherapeutic dose (200 mcg) of IT DMXAA on Day 0 (DMXAA200) or vehicle (NaHCO3). G: Kaplan-Meier survival curves for the experiment in panel S6F, *=p<0.05 by log-rank test. H: Response to DMXAA200 is lost in B16-Vkine tumors implanted in *Batf3*-/- recipients. I: Efficacy of subtherapeutic DMXAA200 in B16-Vkine tumors implanted in *Batf3*-/- recipients is restored following adoptive transfer of iCD103. A subset of B16-bearing tumors did not engraft iCD103, likely attributable to the pauci-immune environment of B16 tumors.

**Supplementary Fig. S7 (related to Fig. 7).**
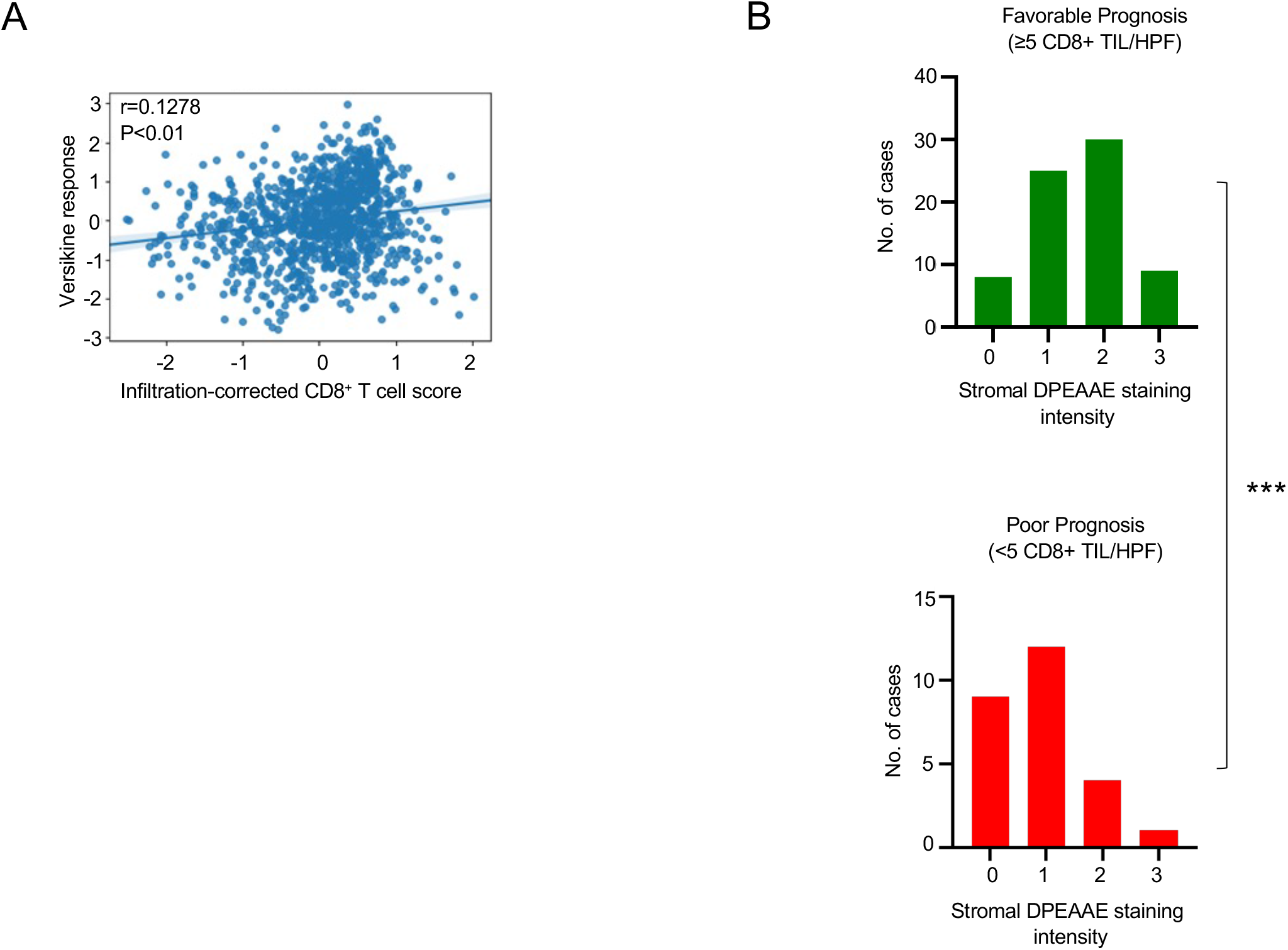
A: Correlation between *in vitro* versikine response signature and corrected CD8+ T cell scores. CD8+ T cell scores corrected for immune infiltration to remove variation associated with immune state. B: Distribution of DPEAAE stromal staining intensity across lung cancer prognostic subgroups [pauci-immune (poor prognosis) and immune-rich (favorable prognosis) at cutoff 5 CD8+ TIL/HPF, p<0.001 by two-tailed Mann-Whitney test].

